# Giant *Starship* elements mobilize accessory genes in fungal genomes

**DOI:** 10.1101/2021.12.13.472469

**Authors:** Emile Gluck-Thaler, Timothy Ralston, Zachary Konkel, Cristhian Grabowski Ocampos, Veena Devi Ganeshan, Anne E. Dorrance, Terry L. Niblack, Corlett W. Wood, Jason C. Slot, Horacio D. Lopez-Nicora, Aaron A. Vogan

**Affiliations:** Department of Biology, University of Pennsylvania, Philadelphia, USA; Department of Plant Pathology, The Ohio State University, Columbus, USA; Laboratory of Evolutionary Genetics, University of Neuchâtel, Neuchâtel, Switzerland; Facultad de Ciencias Agrarias, Universidad Nacional de Asunción, San Lorenzo, Paraguay; Arabidopsis Biological Resource Center, The Ohio State University, Columbus, USA; Ohio Agriculture and Research Development Center, Wooster, USA; Departamento de Producción Agrícola, Universidad San Carlos, Asunción, Paraguay; Systematic Biology, Department of Organismal Biology, University of Uppsala, Uppsala, Sweden

## Abstract

Accessory genes are variably present among members of a species and are a reservoir of adaptive functions. In bacteria, differences in gene distributions among individuals largely result from mobile elements that acquire and disperse accessory genes as cargo. In contrast, the impact of cargo-carrying elements on eukaryotic evolution remains largely unknown. Here, we show that variation in genome content within multiple fungal species is facilitated by *Starships,* a novel group of massive mobile elements that are 110 kb long on average, share conserved components, and carry diverse arrays of accessory genes. We identified hundreds of *Starship-*like regions across every major class of filamentous Ascomycetes, including 28 distinct *Starships* that range from 27-393 kb and last shared a common ancestor ca. 400 mya. Using new long-read assemblies of the plant pathogen *Macrophomina phaseolina*, we characterize 4 additional *Starships* whose past and ongoing activities contribute to standing variation in genome structure and content. One of these elements, *Voyager*, inserts into 5S rDNA and contains a candidate virulence factor whose increasing copy number has contrasting associations with pathogenic and saprophytic growth, suggesting *Voyager*’s activity underlies an ecological trade-off. We propose that *Starships* are eukaryotic analogs of bacterial integrative and conjugative elements based on parallels between their conserved components and may therefore represent the first known agents of active gene transfer in eukaryotes. Our results suggest that *Starships* have shaped the content and structure of fungal genomes for millions of years and reveal a new concerted route for evolution throughout an entire eukaryotic phylum.

## Introduction

A landmark realization of modern evolutionary biology is that genome content is highly variable among individuals of the same species ^1^. Intraspecific genome content variation comprises a species’ pangenome, which consists of core (i.e., consistently present) and accessory (i.e., variably present) orthologs. The pangenome concept has adjusted prior expectations of the rate of adaptive trait emergence and dispersal in populations, as differences in accessory sequence distributions across individuals have been found to arise rapidly and fuel phenotypic divergence ^2–4^. For example, bacterial animal pathogens quickly evolve resistance to antibiotics through the gain of entire genomic regions encoding resistance phenotypes ^5^ and the host range of many fungal plant pathogens often expands through the gain of entire chromosomes encoding host-specific virulence factors ^6^. Identifying the mechanisms generating selectable variation in the distribution of accessory genes among individuals is thus crucial for understanding the genetic bases of adaptation ^7^.

Genome content variation in prokaryotes is primarily driven by mobile elements that in addition to encoding functions for their own movement, also encode accessory genes ^1^. These “cargo-carrying” mobile elements range in size, from smaller transposons to larger genomic islands to entire DNA molecules such as plasmids, and interact with their host genomes in diverse ways ^8–10^. While some mobile elements resemble parasites because their proliferation is predicted to come at the expense of host fitness ^11^, cargo-carrying elements resemble mutualists that cooperate with host genomes because they often carry genes encoding adaptive host traits ^1^. For example, several well known integrative and conjugative elements carry accessory genes mediating advantageous symbiosis and antibiotic resistance phenotypes ^12, 13^. Cargo-carrying mobile elements are frequently transmitted through both horizontal gene transfer (HGT) and vertical inheritance, which together generate extensive genome content variation within and across species ^1^.

In contrast with prokaryotes, the contribution of mobile elements to genome content variation in eukaryotes remains unclear, although is of increasing interest ^14–16^. Eukaryotic genomes are often large and repeat-rich, which complicates the characterization of accessory genes using short-read sequencing methods ^1^. However, long-read sequencing technologies that resolve repetitive sequence content now enable more contiguous genome assemblies in which the movements and associations of mobile elements can be tracked ^14^. Several small cargo-carrying eukaryotic mobile elements in specific species have recently been shown to generate genome content variation by increasing the copy numbers of their accessory genes ^17, 18^. Larger eukaryotic mobile elements have only been identified recently, and though little is known about these elements, initial studies suggest they impact host fitness, that some are horizontally acquirable, and are pervasive across multiple species ^16, 19–22^. Yet despite compelling preliminary evidence, the diversity, phylogenetic relationships, and contributions towards genome and pangenome evolution of large mobile elements are not well understood. Fungi are ideal for investigating these aspects of large mobile element biology due to their relatively small genomes with resolvable repeat content, notable associations between several recently described large mobile elements and phenotypes ^19–, 21^, and a vast diversity of sequenced genomes that enable kingdom-wide analyses.

Here, we investigated the contributions of large cargo-carrying mobile elements to the evolution of Ascomycete fungi, one of the most speciose, ecologically diverse and economically important groups of microbial eukaryotes. We sequenced and searched 12 genomes of the plant pathogen *Macrophomina phaseolina* for evidence of large cargo-carrying mobile elements. We found several giant mobile elements that prompted us to search for similar elements in other Ascomycetes, and test the hypothesis that they all share a common origin. Using *M. phaseolina* as a case study, we then ask how these elements shape the evolution of genome structure, genome content, and ecologically-relevant phenotypes.

## Results and Discussion

### *M. phaseolina* genomes harbor insertions that resemble mobile elements

We sequenced 12 *M. phaseolina* isolates from soybean fields in Ohio, U.S.A. and Paraguay using a combination of Nanopore long-read and Illumina short-read technologies. We used multiple assemblers and gene annotators to obtain near-chromosome level quality draft assemblies and annotations (avg. N50 = 2.43mb; avg. assembly size = 52.31mb)(Figure S1, Tables S1, Table S2, Table S3, Methods). Isolates differed by at most 1% nucleotide identity across all alignable genomic sequence, as was expected given the predominance of clonal reproduction in this species ^23^. In contrast, we found that isolates harbor extensive variation in accessory genes, with gene content diverging 20 times faster than nucleotide sequence per unit of evolutionary time and differing by up to 36% (Figure S2, Table S4). *M. phaseolina*’s pangenome is “open” (Heaps’ alpha = 0.46), meaning that the size of its pangenome is predicted to increase with each newly examined genome. *M. phaseolina*’s pangenome is more open than most other fungal species examined, suggesting that individuals within this species generally harbor high proportions of accessory genes (Figure S2, Table S5, Table S6). We tested whether differences in genome content arise randomly across the genome, or whether they are enriched in specific regions, and found that highly variable accessory genes were significantly enriched in “hotspot” regions (Figure S2, Table S7, Table S8). Accessory gene hotspots do not differ from the overall genome in proportion of SNPs or indels <= 1kb, but are enriched for indels >1kb (Methods), suggesting that gene variability is associated with large structural mutations (Figure S2, Table S9).

We hypothesized that the variation in *M. phaseolina*’s accessory genome and the generation of large structural mutations in hotspots is facilitated by large cargo-carrying mobile elements. We searched the 12 *M. phaseolina* genomes for large insertions that had three key features typified by the recently described giant fungal transposons *Enterprise* and *HEPHAESTUS* ^19, 20^: (1) a predicted gene containing a DUF3435 domain (Protein family accession: PF11917) located at the 5’ end of the element in the 5’-3’ direction. DUF3435 domains are homologous to and share conserved active sites with tyrosine site-specific recombinases that catalyze the excision and integration of bacterial integrative and conjugative elements ^19, 24, 25^; (2) flanking target site duplications (TSDs) that arise when DNA transposons integrate into their target site through staggered double stranded breaks ^26^; (3) terminal inverted repeats (TIRs), which are typically required for the excision of DNA transposons by transposases, although are not present in *Enterprise* ^19, 26^. We found four very large (76,706 - 195,340 bp) elements we call *Voyager, Argo, Phoenix,* and *Defiant* that all have a DUF3435 gene in the required position and often have TSDs and TIRs as well (Figure 1B, Table 1, Table S10). These four elements are discrete and distinct entities that carry diverse arrays of accessory cargo and differ in their target sites and TIRs. However, despite this variability, DUF3435 genes in these elements often co-localize with four additional genes that are present in variable copy numbers and conserved across elements, consisting of DUF3723s, ferric reductases (FREs), patatin-like phosphatases (PLPs) and NOD-like receptors (NLRs) with different functional domains (Table 2, Table S11). We refer to these genes by their predicted function throughout the text.

**Figure 1:**
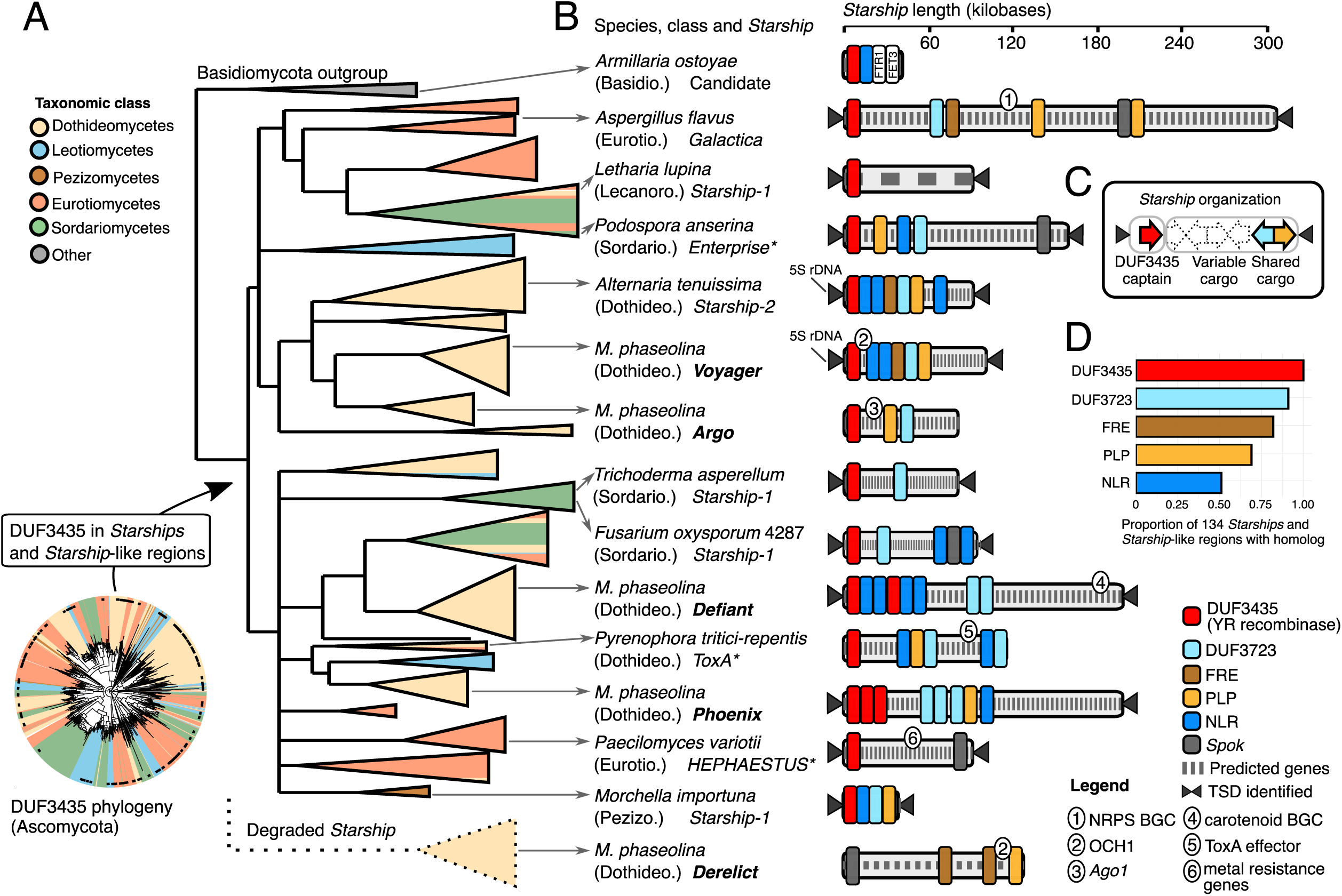
Giant *Starship* elements found across the Ascomycota share conserved components and mobilzie diverse arrays of accessory genes. A) To the bottom-left is a circular maximum likelihood tree of 1890 DUF3435 homologs retrieved from a database containing 12 *Macrophomina phaseolina* genomes generated for this study and 1274 published Ascomycete genomes, where branches are color-coded according to taxonomic class. Black dots along the perimeter indicate DUF3435 sequences found in 194 *Starships* and *Starship*-like regions (60 of which are from *M. phaseolina*), and to the right is a maximum likelihood tree of these sequences and several others of interest, where clades are depicted as triangles for clarity and all branches with <80% SH-aLRT and <95% UFboot support have been collapsed. A clade of degraded *Starships* that lack DUF3435 sequences is juxtaposed below the tree for visualization purposes only. B) Fungal species, taxonomic class, *Starship* name and simplified *Starship* schematics, with approximate locations of genes of interest depicted as colored boxes or numbered. Genes are not drawn to scale. Target site duplications (TSDs), when indicated, were confirmed from alignments of insertion polymorphisms. Names of elements from *M. phaseolina* are in bold, and previously published mobile elements are indicated with an asterisk. C) A schematic showing the general organization of *Starship* mobile elements. D) Proportion of 134 *Starships* and *Starship*-like regions containing homologs to five genes of interest. Gene name abbreviations: FRE (ferric reductase); PLP (patatin-like phospholipase); NLR (Nucleotide Oligomerization Domain-like receptor); *Spok* (spore killer); OCH1 (alpha-1,6-mannosyltransferase); NRPS (non-ribosomal peptide synthetase); *Ago1* (Argonaute). Other abbreviations: BGC (biosynthetic gene cluster). All original trees can be found in the supplementary data files.

**Table 1:** Summary statistics of *Starship* elements in *Macrophomina phaseolina*

| *Starship*<br>element | Avg. length in<br>bp (s.d.) | Avg. number<br>of genes (s.d.) | Avg. % repeat<br>content (s.d.) | Copies per<br>isolate, min-max | Target site |
| --- | --- | --- | --- | --- | --- |
| *Voyager* | 96,567 (13,781) | 37 (9) | 30.4 (10.4) | 0-4 | AGAATACAGGGC |
| *Argo* | 75,148 (14,824) | 26 (4) | 9.5 (2.1) | 0-1 | n.d. |
| *Phoenix* | 74,447 (94,611) | 21 (25) | 44.6 (25.7) | 0-3 | TTTACA |
| *Defiant* | 108,860<br>(57,441) | 28 (15) | 37.7 (15.5) | 0-5 | ACTA |

**Table 2:** Conserved *Starship* cargo genes

| Gene name | *Voyager* reference gene(s) | Protein family domains | Predicted function |
| --- | --- | --- | --- |
| DUF3435 | mp102_11332 | PF11917 (DUF3435) | Tyrosine recombinase |
| DUF3723 | mp102_11356 | PF12520 (DUF3723) | Unknown |
| FRE | mp102_11352 | PF01794 (Ferric_reduc) | Ferric reductase |
|  |  | PF08022 (FAD_binding_8) |  |
|  |  | PF08030 (NAD_binding_6) |  |
| PLP | mp102_11357 | PF01734 (Patatin) | Patatin-like phosphatase |
| NLR | mp102_11337 | PF17107 (SesA) | NOD-like receptor |
|  |  | PF05729 (NACHT) |  |
|  |  | PF12796 (Ank_2) |  |
|  |  | PF00023 (Ank) |  |
|  | mp102_ofg737 | PF06985 (HET) | NOD-like receptor |
|  |  | PF05729 (NACHT) |  |

### *Starships* are a novel group of giant mobile elements widespread across Ascomycetes

We hypothesized that the conserved associations between DUF3435s, DUF3723s, FREs, PLPs, and NLRs are indicative of a “winning” coalition of co-functioning genes maintained through selection ^27^, much like the multi-component translocative machinery of bacterial integrative and conjugative elements ^10^. We therefore predicted that they would appear together in other Fungi. We used the genes in *M. phaseolina*’s elements to query a database of 1274 Ascomycete genomes, and found 134 regions containing homologs to DUF3435 that also colocalized to varying degrees with DUF3723s, FREs, PLPs, and NLRs (Figure 1D, Table S12, Table S13). Using whole genome alignments, we manually annotated the insertion sites, TSDs, and TIRs for 28 of these elements sampled across all extant classes of filamentous Ascomycetes, providing evidence of pervasive and recent mobile activity across one of the most diverse and speciose eukaryotic phyla (Figure 1B, Table S14, Figure S4, Figure S5).

We used the phylogenetic history of DUF3435 genes at the 5’ element ends, which we refer to as “captains”, to characterize the evolutionary relationships between the various elements (Methods). Not only did we find that captains from different elements fall into distinct clades (Figure 1A, Figure S5, Figure S6), but we also found that the tyrosine recombinase domains of all DUF3435 captains group into a monophyletic clade that is as distantly related to bacterial tyrosine recombinases as it is to described tyrosine recombinases found in either fungal *Crypton* DNA transposons ^28^ or eukaryotic retrotransposons, such as DIRS elements ^29^(Figure S7). We conclude that DUF3435-containing genes represent a novel group of eukaryotic tyrosine recombinases that mobilize large amounts of DNA. A study that was simultaneously published with our own independently demonstrated that elements with DUF3435-containing genes are a new monophyletic group of transposable elements (cite V. Kapitonov here). We consider regions bounded by TSDs that contain a DUF3435 sequence at the 5’ end to be a new class of eukaryotic mobile element that we call *Starships* (Figure 1C). Following Vogan et al., 2021, We distinguish between different types of *Starship* elements based on their insertion sites and the phylogenetic relationships between their DUF3435 captains. Regions that contain DUF3435 sequences that co-localize with DUF3723s, FREs, PLPs, and/or NLRs but lack TSDs are referred to as *Starship*-like, and regions that contain at least 3 hits to any of these conserved genes but lack DUF3435 sequence are considered to be degraded *Starships*.

The DUF3435 phylogeny revealed that the captains of different *Starships* likely last shared a common ancestor prior to the divergence of the Pezizomycotina, ca. 400 million years ago ^30^ and thus have a single origin (Figure 1A). This phylogeny unites previously disparate phenomena in fungi, including *Enterprise*^19^*, HEPHAESTUS*^20^, a large mobile region containing the ToxhAT transposon and ToxA effector found in numerous wheat pathogens ^21^, and multiple previously described large structural variants whose origins were unclear (Galactica_Afla, Starship-1_Afum)^31, 32^, by demonstrating that they are all *Starships*. The majority of DUF3435s in the Ascomycota are largely restricted to clades consisting of sequences from the same taxonomic class and multiple class-specific clades exist, consistent with an evolutionary scenario dominated by the vertical inheritance of ancient homologs (Figure S4). This same trend is recapitulated within individual taxa, such as *Macrophomina, Aspergillus* and *Fusarium,* that contain multiple highly diverged and distantly related DUF3435s within their genomes (Figure S5). However, HGT has been implicated in *Starship* transfers among closely related taxa in at least two independent examples ^20, 21^, and rates of HGT are expected to increase with donor-recipient relatedness ^33^, suggesting HGT may be more prevalent at finer grained taxonomic scales beyond the scope of this study.

### *Starship* accessory cargo suggests a range of impacts on host fitness

The diversity of accessory cargo found across *Starships* suggests their interactions with host genomes span the conflict-cooperation spectrum. For example, some predicted cargo suggests mobile element fitness could come at the host’s expense: numerous *Starship*s contain homologs to spore killing (*Spok*) genes that encode a toxin-antitoxin meiotic drive system capable of killing developing sexual progeny in *Enterprise* ^19, 34^ (Figure 1B, Table S13). In contrast, many cargo genes have putatively adaptive functions that would hypothetically benefit the host in certain environments: several carry biosynthetic gene clusters (BGCs) involved in the production of secondary metabolites (e.g., *Defiant* and *Galactica*); others carry characterized virulence factors or homologs of known virulence factors (e.g., *ToxA* ^21^, Ago1 in *Argo* ^35^, OCH1 in *Voyager and Derelict* ^36–38);^ *HEPHAESTUS* contains several genes characterized to contribute to heavy metal resistance^20^; finally, FREs that are conserved across multiple *Starships* are part of a large metalloreductase family with several characterized members, suggesting some confer benefit to the host, for example, in the acquisition of iron in growth limiting environments or contributions to pathogen virulence ^39–41^. While the few *Starship*-like regions we identified in our Basidiomycete outgroup lacked FRE homologs, they contain homologs to Fet3p and Ftr1p which form a high affinity iron permease complex in yeast ^42^, suggesting a functional association convergently evolved between iron metabolism and *Starships* (Figure 1B). Carrying host-beneficial cargo may be an especially important trait for elements that are predominantly inherited through vertical means, as their own fitness is closely aligned with their host’s reproductive success ^43^.

*Starship* activity shapes the distribution of accessory genes in *M. phaseolina* populations

We carried out an in-depth characterization of *M. phaseolina*’s *Starships* to better understand how these larger elements shape variation in accessory genome content. *Starship* elements in *M. phaseolina* differ in copy number, genetic cargo, and generally have little alignable sequence in common such that they each contribute in distinct ways to content divergence across *M. phaseolina* isolates (Figure 2A, Table 1, Table S15). Extensive content variation even exists within certain *Starship* element types, such as *Phoenix* and *Defiant*. For example, we found a putative cargo turn-over event in two *Defiant* elements that carry either a carotenoid BGC or a non-ribosomal peptide synthetase (NRPS) BGC (Figure 2D). Within any given genome, 85-100% of *Starships* overlap with accessory gene hotspots indicating that they are hotspots for gene gain and loss (Methods, Figure 2C, Table S16). Specific *Starship* elements are further linked to the expansion of accessory gene families they carry as cargo (Figure 2B). For example, over half of the OCH1 homologs (59/109) and just over a third of the FRE homologs (112/316) in the *M. phaseolina* population are located in *Starship* or *Starship*-like regions (Figure 2B, Figure S8, Figure S9). Conserved cargo genes and functional domains carried on *Starships*, including OCH1, PLP, FRE, DUF3723 and the HET and NACHT domains in NLRs, are typically restricted to clades of *Starship*-specific sequences, indicative of cargo-*Starship* co-diversification (Figure 2B, Figure S8, Figure S9, Figure S10, Figure S11, Figure S12, Figure S13). Content diversity and flexibility, coupled with a transposition mechanism, underscores the potential for *Starships* to shape the distribution of accessory genes among individuals.

**Figure 2:**
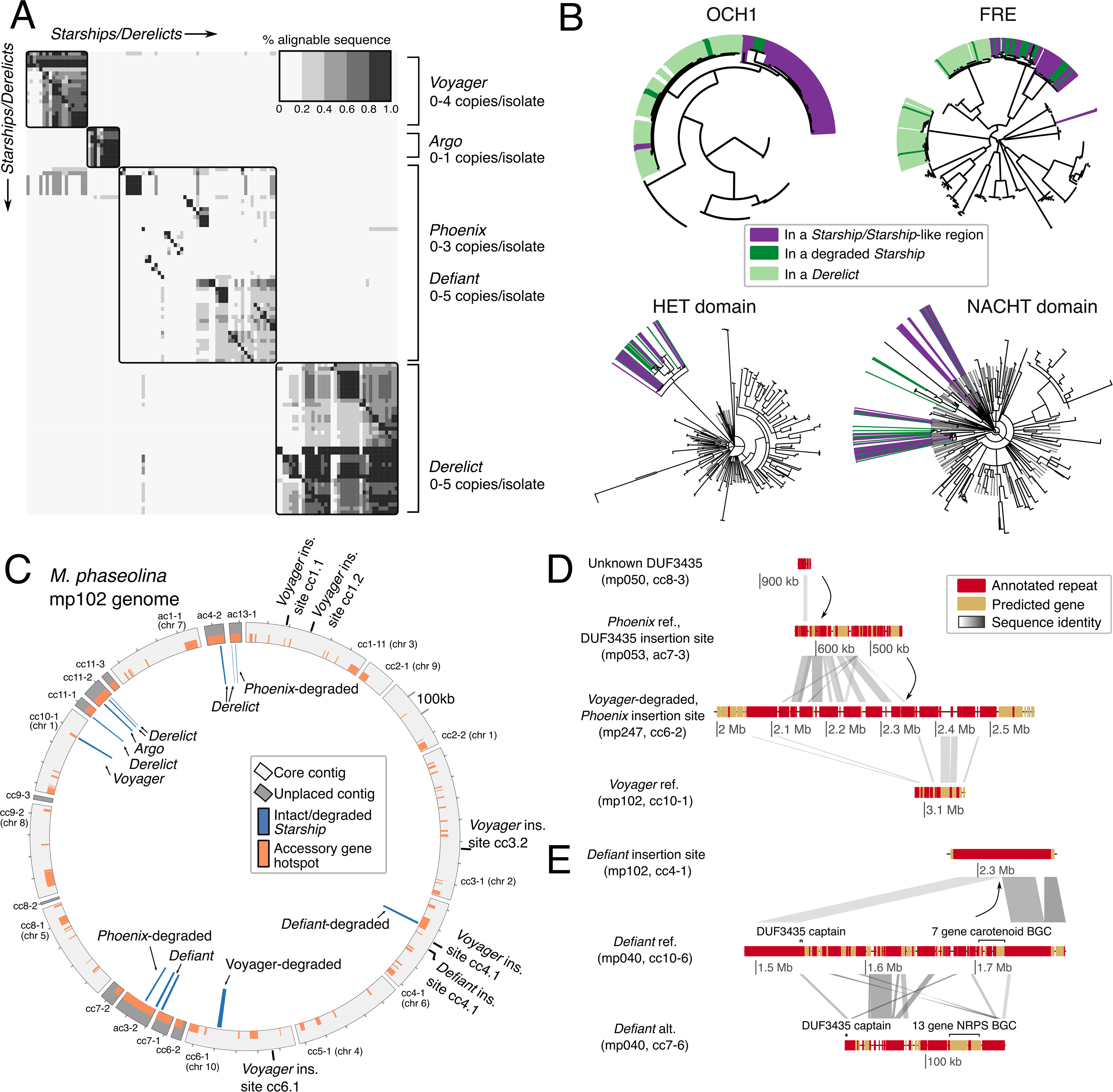
Distinct *Starship* elements in *Macrophomina phaseolina* vary in copy number across individuals and generate intraspecific variation in genome content. A) A heatmap summarizing the percentage of pairwise alignable sequence between all *Starships* and *Starship-*like regions, as well as the degraded *Starship* region *Derelict*, found across 12 *M. phaseolina* isolates, where rows and columns represent individual regions arranged by hierarchical clustering according to similarity in their alignment scores. B) Maximum likelihood phylogenies of genes homologous to OCH1 and FREs, as well as genes containing and HET and NACHT domains found in fungal NLRs, retrieved from the *M. phaseolina* genomes, where all branches with <80% SH-aLRT and <95% UFboot support have been collapsed. Genes found in regions of interest are highlighted by different colors. C) A circos plot depicting all contigs >100kb in the *M. phaseolina* mp102 genome. Core contigs are partially alignable across all isolates, while unplaced contigs are not. Regions of interest are labeled and visualized on the inner track. Sites with resolved *Starship* insertions in other genomes, but absent in mp102, are labeled on the outside track. Accessory gene hotspots are labeled in orange on the outer track. D) A schematic depicting a sequential *Starship* insertion event where an unknown DUF3435 gene is found nested in *Phoenix*, which itself is nested in a degraded *Voyager* element in isolate mp247 on contig cc6-2. E) A schematic depicting a cargo turn-over event between two *Defiants* carrying either a carotenoid or non-ribosomal peptide synthetase (NRPS) biosynthetic gene cluster (BGC). Gene name abbreviations: OCH1 (alpha-1,6-mannosyltransferase); FRE (ferric reductase); NLR (Nucleotide Oligomerization Domain-like receptor);

Not only do *Starship* elements actively generate new variation, but remnants of degraded *Starship*-like regions found throughout the genomes of *M. phaseolina* suggest their impact extends beyond the life of the element. For example, we discovered a group of regions, which we call *Derelict,* that have *Starship*-specific cargo but lack DUF3435 homologs (Figure 1A, Table S11). Like active *Starships, Derelicts* show large swaths of similarity among isolates and copies (avg. 41% alignable sequence), but otherwise are distinct from other elements (Figure 2A). They have homologs of FRE, DUF3723, and OCH1 that form large monophyletic clades within their phylogenies, and these sequences are in general more closely related to their homologs in other *Starships* than they are to non-*Starship* associated sequences, suggesting that *Derelicts* are old copies of a *Starship* whose captain degraded but whose cargo has persisted (Figure 2B, Figure S9, Figure S8, Figure S13). We find many additional *Starship*-like regions that similarly lack DUF3435 homologs but have conserved *Starship* content and occasionally aligned to characterized *Starship* elements, suggesting they have recently degraded (Table S11). Across all genomes, 72-100% of these degraded regions overlapped with an accessory gene hotspot, indicating that they have remained a source of genome content variation among individuals (Table S16). Across the 1274 Ascomycete genome database, we similarly found 102 *Starship*-like regions that did not have DUF3435 genes, but had at least four hits to FRE, DUF3723, NLRs, and PLP sequences, suggesting these regions may also be degraded *Starship* elements (Table S13). Evidence of cargo that persists after element degradation implicates *Starship* birth-and-death cycles as a fundamental mechanism shaping variation in genome content among individuals.

*Starships* contribute to standing variation in the structure of *M. phaseolina*’s genome

Many fungal genomes are compartmentalized into accessory chromosomes that are variably present among individuals, and core chromosomes with large tracts of conserved synteny across individuals ^44^. In *M. phaseolina*, we identified 10 large “core contigs” that are largely syntenic, often have telomeric repeats at either end, and are present across all individuals (Figure 3, Figure S3, Table S17, Table S18, Methods). We found that on average, 88.6% of each *M. phaseolina* genome is captured by these core contigs, while the rest is located in unplaced contigs that do not align to the core contigs (Table S19). However, core contigs in the *M. phaseolina* population are highly rearranged and contain large tracts of variably present sequence, such that none of the core contigs in the eight highest quality assemblies share the same structure or content across all isolates (Figure S3), consistent with previous observations in this species ^45^.

**Figure 3:**
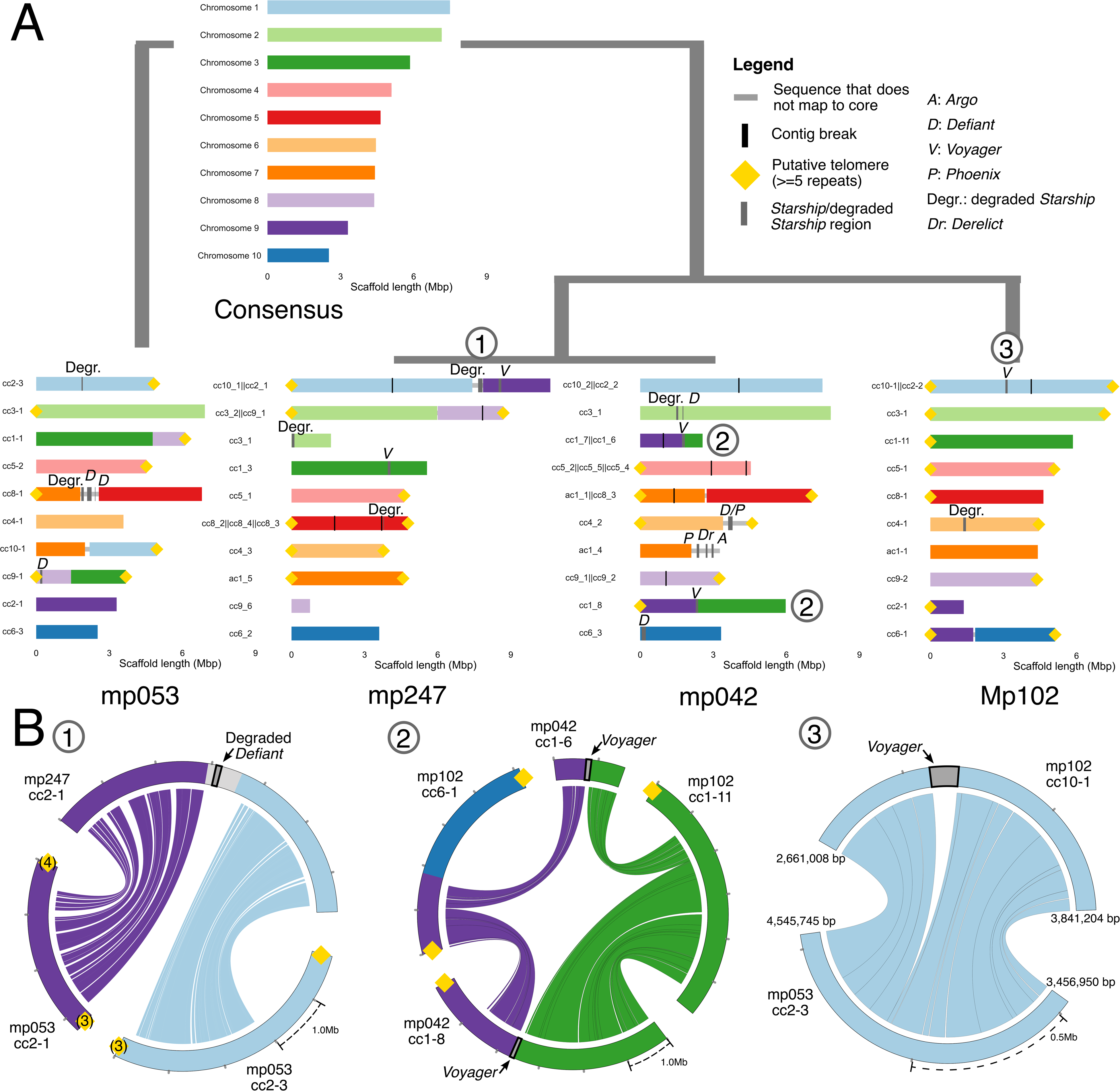
Several *Starship* elements are associated with large structural variants across *Macrophomina phaseolina* genomes. A) Alignment-based chromosome maps of four *M. phaseolina* isolates and a hypothetical ancestral genome, where contigs that share homology are colored and ordered according to their putative chromosome of origin. Putative telomeres are labeled at contigs ends, and consist of at least 5 sequential telomeric repeats, unless indicated otherwise by bracketed numbers. Unplaced contigs are not illustrated. B) From left to right, circos-based nucmer alignments of (1) a putative chromosomal fusion, (2) a reciprocal translocation, and (3) a *Voyager* insertion polymorphism. Contigs are labeled according to chromosome of origin, or otherwise colored gray. *Starships* and degraded *Starships* are drawn in dark gray, labeled and outlined in black. Labeled numbers correspond to events depicted in both the chromosome maps and the circos alignments.

We therefore examined associations between *M. phaseolina*’s *Starships* and large-scale differences in genome structure, which may be facilitated by their ability to insert into otherwise conserved sequence tracts (Figure 3B, event 3). We identified 37 large translocations, inversions and chromosome fusions in the *M. phaseolina* population, and 30% of these had *Starships, Starship-*like regions, or degraded *Starships* overlapping or near their breakpoints (Table S20). For example, a degraded *Defiant* element is found at the junction of a putative fusion between core chromosomes 9 and 1 in *M. phaseolina* isolate mp247, and a reciprocal translocation between core chromosomes 9 and 3 in *M. phaseolina* isolate mp042 occurs precisely at a *Voyager* insertion (Figure 3B, events 1 and 2). Due to their sheer size, sequence similarity, and the fact that they are themselves colonized by multi-copy mobile elements including LTRs (Table S10), *Starship* insertions may provide opportunities for rearrangements to occur through ectopic recombination ^46^, although our data cannot confirm whether they are the causal mechanism. However, 95% of the rearrangements overlapped with some sort of mobile element, in general agreement with previous observations that mobile elements are associated with large scale rearrangements in fungal genomes ^46, 47^.

*Starship* activity is also linked to the active growth of large genomic regions whose presence varies among individuals. Gene phylogenies of DUF3435s and other conserved cargo suggest *Starships* can sequentially nest within each other, generating large tracts of accessory sequence (Figure S6). For example, a degraded *Voyager* insertion in *M. phaseolina* isolate mp247 has at least three *Starship* or *Starship-*like elements that have sequentially inserted into each other to produce a repeat-rich region spanning 221,058 bp (Figure 2D). Mobile element nesting is a common mechanism of bacterial integrative and conjugative element growth ^48^ and has also been observed in *HEPHAESTUS* ^20^. In addition to generating variable regions on otherwise core contigs, *Starships* and *Starship*-like regions may also contribute to the birth and/or growth of accessory chromosomes. Contig ac11-1, a putative accessory chromosome in *M. phaseolina* isolate mp053, is ∼ 800 kb, with *Starship-like* regions composing roughly 16% of its length.

### *Voyager*: a massive mobile element targeting 5S rDNA

The most active *Starship* in the *M. phaseolina* population is *Voyager,* ranging in copy number from 0-4 across individuals (Figure 4A). We resolved 6 different *Voyager* insertion sites that have clear breakpoints and high Illumina read coverage across occupied and empty sites (Figure 4C, Table S21). In isolates with a given *Voyager* insertion, flanking regions are syntenic with the corresponding insertion site in isolates that do not have the insertion, ruling out the alternative explanation that *Voyager* has moved by chromosomal translocation. Based on these resolved insertion sites, we recovered 15 full-length copies of *Voyager* with identifiable TSDs and TIRs (Figure 4B, Table 1). We also found an additional 7 fragmented copies from the two lowest quality assemblies (mp124 and mp194) that had recoverable DUF3435 sequences, but were excluded from the following summary statistic calculations. *Voyager* elements range from 74,275 bp to 124,223 bp in length (avg. 96,567 bp), contain between 10-47 predicted protein-coding sequences (avg. 37), and are alignable from start to end. Variation in size and in protein-coding sequences among *Voyager* copies is largely due to variation in transposable element content, which ranges from 15-50% (avg. 31.2%; Table S21).

**Figure 4:**
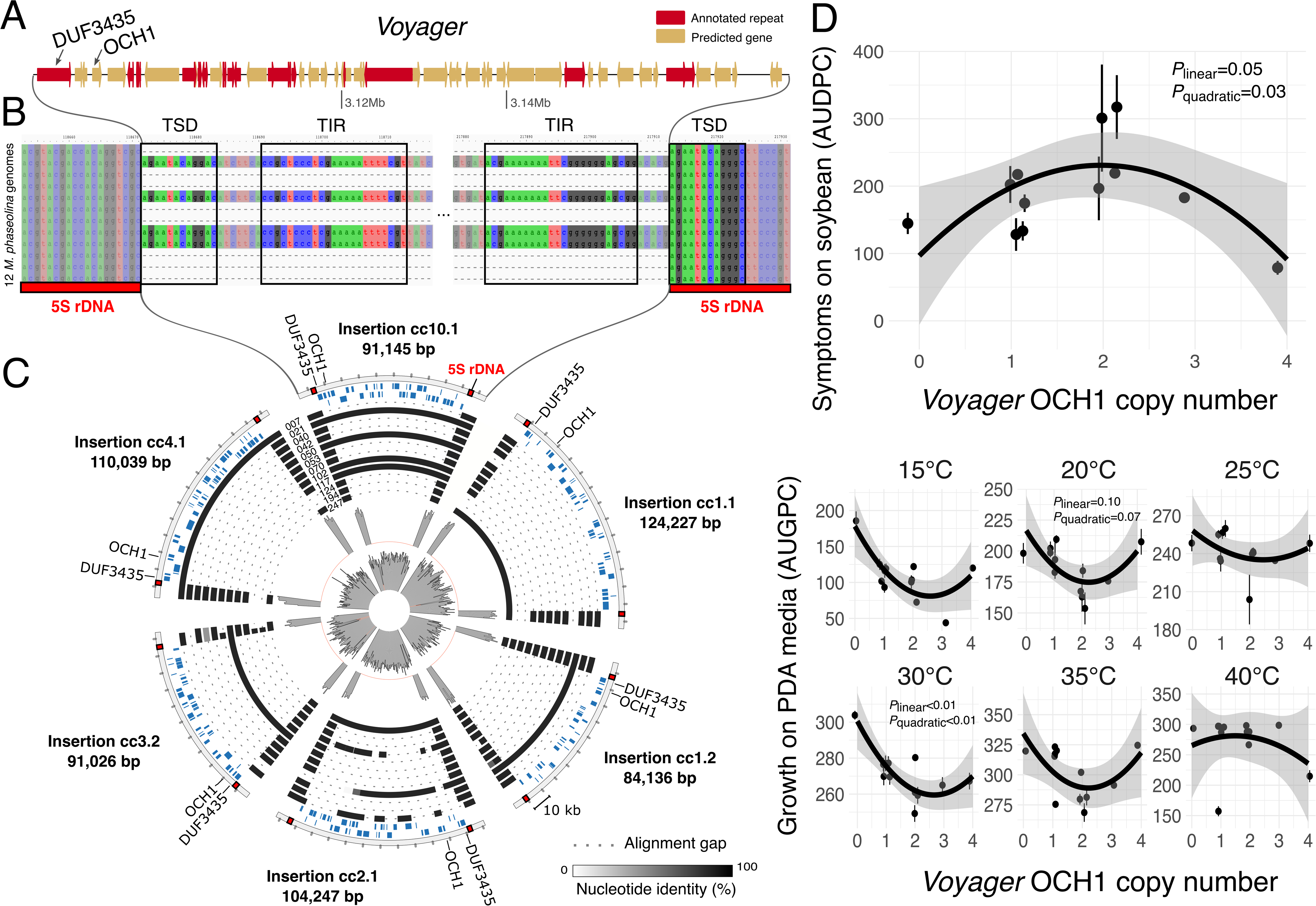
Increasing *Voyager* OCH1 copy number has contrasting effects on *Macrophomina phaseolina*’s pathogenic and saprotrophic growth. A) A schematic of the reference *Voyager* element in *M. phaseolina* mp102 (ID: mp102|voyager|h11) depicting all predicted genes and annotated repetitive elements B) A multiple sequence alignment of the *Voyager* insertion cc10.1, with annotated target site duplications (TSDs) and terminal inverted repeats (TIRs) C) A circos plot of six fully-resolved *Voyager* insertions with 10kb flanking regions, labeled by insertion ID and length. Tracks 1-2 depict genes in the + and - orientation, respectively. Tracks 3-14 depict the percent nucleotide identity of sequence alignments at each locus across 12 *M. phaseolina* genomes, arranged by increasing isolate number. The density plot in Track 15 depicts median Illumina coverage across intact 5S rDNA insertion sites in isolates with empty insertion sites, and ranges from 0-77x. The density plot in Track 16 depicts median Illumina coverage across *Voyager* insertions and their flanking regions, and ranges from 0-100x. D) Scatterplots (jitter = 0.05) depicting mean area under the disease progress curve (AUDPC; top) and mean area under the growth progress curve (AUGPC; bottom) phenotypes as a function of the number of OCH1 homologs found in *Voyager Starships,* for each of the 12 isolates. AUGPC was measured at 6 different temperatures. Error bars indicate +/- standard error, and are visible if they exceed the area of the point. Predicted quadratic equations with shaded 95% confidence intervals are superimposed on the plots. *P*values for significant (*P* <0.05) and marginally significant (*P<*0.10) linear and quadratic coefficients are shown, and others in Table S24. Abbreviations: OCH1 (alpha-1,6-mannosyltransferase); PDA (potato dextrose agar).

Although *Voyager* copies are found at multiple sites in any given genome, they are always inserted into a copy of the 5S rDNA gene, 20 bp from the gene’s 5’ start site and 110 bp from the 3’ end. The insertion of *Voyager* into the 5S rDNA gene shifts the free energy of the 5S rRNA’s predicted secondary structure from an estimated -42.22 kcal/mol to -38.49 kcal/mol such that we predict it loses function. Due to experiencing strong purifying selection^49^, rDNA sequences have been proposed as stable mobile element target sites enabling long-term persistence^50^. Similarly, many bacterial integrative and conjugative elements insert in or near tRNA genes which would also provide stable target sites ^9, 24^. We estimate that *M. phaseolina* has upwards of 45 copies of 5S rDNA genes dispersed across its genome that are fixed in the population, similar to copy numbers observed in other Ascomycete fungi^49^. *M. phaseolina* has at least 2 distinct variants of the 5S rDNA gene, and *Voyager* insertions and target sites are only present in one of them, raising the possibility that changes to the 5S sequence of this species are selected to avoid being disrupted by *Voyager*. Additionally, we identified a *Starship* in *Alternaria tenuissima* that has distinct cargo but whose captain is closely related to *Voyager*’s and whose target site is similarly located inside 5S rDNA sequence (Figure 1B). This suggests that targeting conserved host genes is a broadly successful *Starship* strategy even though it may open the door to potential conflict with host genomes.

### *Voyager* has contrasting associations with pathogenic and saprophytic growth

Insight into the genetic bases of virulence in *M. phaseolina* is critically needed because this pathogen represents an emerging threat to food security: it can infect more than 500 plant species, including many agricultural crops, and is increasingly expanding into new geographic regions. Although mobile element insertions in rDNA tend to negatively impact growth ^51, 52^, the majority of *Voyager* elements also carry homologs of the virulence factor OCH1, a conserved α-1,6-mannosyltransferase that contributes to cell wall integrity and pathogenicity in diverse plant and animal pathogens ^36–38^. We therefore hypothesized that *M. phaseolina*’s growth on its host would increase with the number of OCH1-carrying *Voyager* elements present in the genome. We inoculated soybean seedlings in a greenhouse, and measured the area under the disease progress curve (AUDPC) as a proxy for pathogenic growth (n = 12 isolates, 12 replicates/isolate; Table S22). We similarly inoculated potato dextrose agar (PDA) plates placed at six different temperatures in an incubator, and measured the area under the growth progress curves (AUGPC) as proxies for saprophytic growth (n = 12 isolates, 3 replicates/isolate/temperature; Table S23). Using linear models corrected for phylogenetic relatedness among isolates (Table S24), we found a significant positive linear and negative quadratic relationship between *Voyager* OCH1 copy number and pathogenic growth (y = 138.3 + 93.6x - 22.5x^2^; *P*_linear_=0.05, *P*_quadratic_=0.03) that suggests the predicted contributions of *Voyager* OCH1 to pathogenicity are greatest at intermediate copy number but then decline. Conversely, we found a marginally significant negative linear and positive quadratic relationship for saprophytic growth at 20°C (y = 201.1 -31.6x + 7.3x^2^; *P*_linear_=0.10, *P*_quadratic_=0.07) and a significant negative linear and positive quadratic relationship for saprophytic growth at 30 °C (y = 306.7 - 35.9x + 7x^2^; *P*_linear_<0.01, *P*_quadratic_<0.01) that together suggest the contributions of *Voyager* OCH1 to saprotrophy are lowest at intermediate copy number. These results are consistent with a scenario where *Voyager* incurs an ecological trade-off between *M. phaseolina*’s ability to grow on or off of its host, and suggests alternating host and non-host environments help maintain variation in virulence phenotypes in natural populations. However, our small sample size (n=12) demands that observed trends be confirmed once more individuals are genotyped for *Voyager* insertions. The complex associations of *Voygager* with fungal-plant interactions may predate the origin of the *Macrophomina* genus because we identified a putatively vertically inherited copy of *Voyager* in the closely related plant pathogen *Botryosphaeria dothidea* that has the same target site and also carries OCH1, but may not be presently active due to an LTR insertion into the reading frame of the captain (Table S14, Figure S8). Vertical inheritance of the same element strengthens the hypothesis that conserved host genes provide stable target sites, and underscores how this strategy may enable persistence through speciation events.

### Are *Starships* agents of active gene transfer in eukaryotes?

*Starships* are distinct from other eukaryotic mobile elements not only in their consistently large size, but in that many of them share a conserved set of accessory genes (Table 2). Although the functions of these genes are not known, they present parallels to translocative and dispersive modules in bacterial integrative and conjugative elements, especially those found in Actinomycetes who produce coenocytic hyphal networks resembling filamentous Ascomycete fungi. Like many of these bacterial elements, *Starships* are likely mobilized by site-specific tyrosine recombinases located at the 5’ end of the element ^10, 53^. DUF3723s are homologous to DNA-binding bacterial chromosome segregation genes (TIGRFAM accession: TIGR02168), and present a functional analogy to DNA translocases that catalyze bacterial element transfer ^10^. NLRs with HET, NACHT and PLP domains govern hyphal fusion events in fungi ^54, 55^ and present parallels with Nudix hydrolase genes that govern Actinomycete hyphal fusion prior to integrative and conjugative element transfer ^10^. Genes with NUDIX domains are also found in several *Starships* (*Galactica, Starship-2_Afla*) (Table S13). Together, these shared features suggest *Starships* encode mechanisms not only for their own excision and integration but for their transfer between individual genomes, and provide testable hypotheses for elucidating a mechanism of fungal HGT, which has long been searched for.

## Conclusion

### *Starships*: new frontiers for eukaryotic genome evolution

*Starships* represent a dramatic shift in our understanding of how mobile elements shape eukaryotic genome and pangenome evolution ^14^. They add to the growing realization that eukaryotic genomes, ranging from fungi to algae to animals, are frequently colonized by large mobile elements, and their activity casts the impact of these other elements in a new light ^14–16^. Using the plant pathogen *M. phaseolina* as a model, we demonstrate that *Starships* link the processes of mobile element activity, structural variant generation and genome content evolution such that *Starship* birth-and-death lifecycles likely hold multi-faceted consequences for their hosts. Our work with *M. phaseolina* further suggests that *Starship* characterization will be important for understanding the emergence of virulence in pathogen populations. *Starships* carrying diverse accessory genes are found in fungi ranging from saprotrophs to lichens to human and plant pathogens and are further known to encode adaptive traits ^20, 21^, which coupled with their potential role in mediating life history trade-offs, underscores their vast potential for shaping ecological outcomes across an entire kingdom. The *Starship* activity that we observe in present-day fungal populations likely represents only a fraction of their historical impact, as their activity has been ongoing for millions of years. Together, our results establish a novel framework for investigating how mobile element cargo carriers shape the (pan)genomic bases of fungal adaptation, pushing our understanding of eukaryotic evolution to bold, new frontiers.

## Supplemental information

### Supplemental Tables

Table S1: Macrophomina phaseolina isolate metadata

Table S2: M. phaseolina genome assembly quality assessment

Table S3: M. phaseolina contigIDs and corresponding accessions

Table S4: All pairwise M. phaseolina genome comparison statistics (used for Figure S2A)

Table S5: Metadata of fungal genomes used for pangenome analyses (used for Figure S2B)

Table S6: Pangenome rarefactions (used for Figure S2B)

Table S7: M. phaseolina accessory gene hotspot coordinates

Table S8: Accessory gene hotspot summary statistics

Table S9: Mutation densitions in accessory gene hotspots and non-hotspot regions

Table S10: Starship and Starship-like region summary statistics (summarized from Table S11)

Table S11: Coordinates and annotations of all genes in all Starship regions in M. phaseolina

Table S12: All metadata of additional genomes that were searched for Starships

Table S13: Coordinates and annotations of all genes in all identified Starship and Starship-like regions

Table S14: Metadata of all annotated Starships identified in this study

Figure S15: All pairwise nucmer alignments between Starships and Starship-like regions in M. phaseolina

Table S16: Per M. phaseolina genome Starships and Starship-like regions summary statistics (summarized from Table S11)

Table S17: All contigs that align to core M. phaseolina contigs (putative chromosomes)

Table S18: Telomeric repeats in Nanopore reads mapping to contig ends

Table S19: Summary statistics of core and unplaced contigs across all M. phaseolina isolates

Table S20: All coordinates and read coverage of large structural variants

Table S21: All coordinates of identified Voyager insertion sites plus 10kb flanking regions

Table S22: Area under the disease progress curve (AUDPC) measurements on soybean seedlings

Table S23: Area under the growth progress curve (AUGPC) measurements on PDA media

Table S24: Phylogenetic generalized least squares models and test statistics

Table S25: Sequence information gained by assembly merging

Table S26: RIPper output

Table S27: Mating-type loci in M. phaseolina

### Supplemental Figures

Figure S1: genome overview

Figure S2: pangenome overview and hotspots

Figure S3: chromosome maps

Figure S4: DUF3435mycoDB

Figure S5: starshipsMycodb

Figure S6: DUF3435macph

Figure S7: YRtree

Figure S8: OCH1macph

Figure S9: FREmacph

Figure S10: HETmacph

Figure S11: NACHTmacph

Figure S12: PLPmacph

Figure S13: DUF3723macph

Figure S14: methodsOverview

#### Data availability

This Whole Genome Shotgun project has been deposited at DDBJ/ENA/GenBank under the Biosample accessions SAMN17156263-SAMN17156274. All additional supporting data, including predicted protein sequences, GFFs, repeat libraries, characterized *Starship* elements, sequence alignments and newick tree files are available from the following Figshare repository (DOI: 10.6084/m9.figshare.17185880).

## Acknowledgements

We would like to thank Megan McDonald, Hannah Wilson, Alexander Idnurm and Andrew Urquhart for inspiring discussions on large mobile elements; S. Lorena Ament-Velásquez, for the development of scripts that were invaluable to the outcomes of this research; Paula Moolhuijzen, who supplied several genomes for the pangenome analysis; Thomas Badet, who supplied scripts for genome alignment visualization and provided helpful feedback on the manuscript; the following funding agencies for their support: Formas for the grant 2019-01227 (to AAV) and the Ohio Soybean Council for Project 17-R-13 (to TLN and HDLN).

## Contributions

EGT, HDLN and AAV conceptualized the research. EGT and AAV conducted formal analyses, with contributions from HDLN and TR. ZK, CGO and VDG provided materials. HLN secured research grant funding, and JCS, CWW, AED and TLN provided additional personnel support. EGT and AAV wrote the original draft, while EGT, AAV, JCS, CWW and HDLN participated in reviewing and editing the manuscript.

## Methods

### Macrophomina phaseolina isolation

Composite soil samples were collected from six soybean fields in Paraguay (2014-2015) and six in Ohio, USA (2013-2014)(Figure S1). A 10-hectare area was defined in each field and with a 2.5-cm-diameter cylindrical probe, 20 to 25 soil cores were collected at a 25 cm depth and thoroughly mixed. From each composite soil sample, *M. phaseolina* was isolated from a 1-gram subsample of air-dried soil as previously described^56^. Colony forming units (CFU) morphologically identified as *M. phaseolina* were transferred to a new PDA plate to obtain pure isolates.

### Phenotyping on soybean seedlings

A modified soybean cut-stem inoculation technique was used to assess the pathogenicity of the *M. phaseolina* isolates^57, 58^. In the greenhouse, soybean plants were grown to the second stage of vegetative growth^59^. The stem was cut 25 mm above the second unifoliate node with a surface sterilized scalpel. A mycelial plug from each *M. phaseolina* isolate was placed on top of the cut stem. Control plants were prepared by placing an uninoculated PDA plug on the cut stem. A 200 µl pipette tip was used to cover the stem and mycelial plug to ensure contact of the fungus and the plant and to prevent the plug from drying out. Plants were arranged in a randomized complete block design with 6 blocks and two plants per experimental unit. Soybean seeds were sown in 48-cell 4×12 plastic trays filled with potting mix (ProMix BX, Premier Horticulture Ltd.) and thinned to one plant per cell post emergence. Two adjacent plants were considered an experimental unit, and one block was composed of a flat containing 26 plants (used for 12 *M. phaseolina* isolates and 1 control). A total of six blocks were placed on a greenhouse bench and covered with a plastic tent, to maintain temperature and humidity. Plants were watered as needed, temperature maintained at 27°C ± 3°C, and a 16-h photoperiod. Three days post inoculation (dpi), pipette tips were removed, and the length of the necrotic lesion was measured at 3, 6, 9, and 12 dpi. The area under the disease progress curve (AUDPC) was calculated using the agricolae package in R (version 4.1.0) for each isolate in each block^60, 61^. We replicated the above experiment in 2018 and in 2021. Analysis of variance revealed that the two experiments were not statistically different (*P* = 0.532), therefore, data from both experiments was pooled. An average AUDPC for each isolate was then determined by averaging all values across the 12 blocks from the two experiments.

### Phenotyping on PDA media

From each *M. phaseolina* isolate, a 6-mm-diameter mycelial plug was placed in the center of a 100-mm-diameter Petri dish containing PDA. Each Petri dish was considered an experimental unit and arranged in a randomized complete block design with 3 blocks, each corresponding to a different shelf in an incubator. Experimental units were incubated in the dark in the incubator for each of the above temperatures. Colony diameter was measured every 24 hours for five days and these measurements were used to calculate the area under the growing progress curve (AUGPC) for each isolate in each block using the agricolae package in R (version 4.1.0) as previously described^60, 62^. An average AUGPC for each isolate was determined by averaging values across the 3 blocks. This experiment was repeated using incubators set at 15, 20, 25, 30, 35, and 40°C

### DNA extraction, library prep and genome sequencing

Isolates were grown in potato dextrose broth (PDB) by shaking cultures in the dark for 3-5 days at 27°C. Mycelia were sieved through autoclaved Miracloth (EMD Millipore, Darmstadt, Germany) and rinsed with sterile distilled water. Mycelia were flash-frozen in liquid nitrogen and stored at -20°C until genomic DNA extraction. Frozen fungal mycelia were ground with a mortar and pestle and genomic DNA was extracted using the DNeasy Plant Mini Kit (Qiagen, GmbH, Germany) following manufacturer’s protocol. DNA libraries were prepared for short-read Illumina Paired End 300 sequencing using the Indexed Truseq DNA Library Preparation Kit (cat.#: FC-121-2001). Four samples were sequenced per Mi-Seq Illumina flow cell with 600 cycles (Illumina, San Diego, CA, USA). DNA libraries for isolates mp007, mp021, mp124, and mp194 were prepared for Oxford Nanopore long read sequencing using the indexed Nanopore Ligation Sequencing Kit SQK-LS108, while all others were prepared using SQK-LS109. Two DNA samples were sequenced per MinION FLO-MIN-106 flow cell with R9.4.1 1D chemistry (Oxford Nanopore Technologies, New York, NY, USA). All library preparation and sequencing was carried out by the Molecular and Cellular and Imaging Center (Wooster, OH, USA).

### Read processing, genome assembly, and chromosome annotations

Illumina reads were trimmed and quality filtered using Trimmomatic v0.36 (ILLUMINACLIP:TruSeq3-PE.fa:2:30:10:11 HEADCROP:10 CROP:285 SLIDINGWINDOW:50:25 MINLEN:100). Nanopore reads were basecalled and de-multiplexed with Albacore v2.2.4 (default settings; Oxford Nanopore Technologies), and filtered with Porechop v0.2.3 to remove adaptors, to only retain reads with at least 1 barcode, and to split chimeric reads (default settings; https://github.com/rrwick/Porechop). Given that different genome assemblers often include different amounts of sequence in their final output, we merged the non-redundant output of multiple assemblers to obtain the most complete assemblies possible. First, filtered Illumina and Nanopore reads were used to generate hybrid assemblies with SPAdes v3.12 (-- careful, -k 55,77,99,127 --cov-cutoff auto)^63^. Then, unfiltered Illumina and filtered Nanopore reads were used to generate hybrid assemblies with MaSuRCA v3.2.8 (default settings)^64^ that underwent 10 rounds of polishing by Pilon v1.22 using filtered Illumina reads to correct bases, fix mis-assemblies, and fill gaps (default settings)^65^. All contigs from the SPAdes assembly were then aligned to the MaSuRCA assembly using nucmer v4.0.0beta (delta-filter options: -q)^66^, and contigs fully unique to the SPAdes assembly were added to the MaSuRCA assembly to obtain the final assemblies, which were modestly larger than the MaSuRCA assemblies alone (Table S25). We calculated pairwise percent nucleotide identity of alignable sequence between all assemblies as “Length of alignable and identical sequence / Total length of alignable sequence” using nucmer v4.0.0beta (--maxmatch).

We classified contigs into provisional core chromosomes using multiple criteria. First, to leverage population-level data that could help in the annotation of telomeres from specific isolates, we mapped all long-read sequences to the ends of each contig >10kb in each isolate. We kept all reads that aligned to at least 1000bp at the contig ends, and considered at least 5 sequential fungal telomeric repeats (TTAGGG)n to indicate the presence of a putative telomere. We then grouped contigs into homologous sets using a protocol inspired by graph-based methods to derive gene ortholog groups^67^. We conducted all possible pairwise genome alignments using the minimap2 aligner in RagTag^68^, which outputs a grouping confidence score for all contig pairs calculated as the number of base pairs a query contig covers in its assigned reference contigs divided by the total number of covered base pairs in the entire reference genome. We took the best confidence score >0.5 for each contig pair, and input them into the MCL clustering algorithm(inflation = 1.7)^69^ such that sets of contigs with similar scores were designated as homologous. However, this alignment-based method failed to account for large-scale rearrangements and translocations, such that contigs that were clearly rearranged chromosomes were being grouped together. We therefore took a conservative approach for our final classification of contigs into core chromosomes by aligning the two highest-quality genomes that were most distantly related to each other (mp102 and mp053; Figure S1), and designating all of their shared chromosome-scale contigs as ‘core’ (Figure S3). We then re-aligned contigs >1kb in all genomes to mp102’s core chromosomes using the nucmer aligner in RagTag (-i 0.2 -a 0.2 -f 1000 --remove-small --nucmer-params ’--mum -l 200 -c 500’), and designated all contigs aligning to these core chromosomes as ‘core’, and all others as ‘unplaced’. To generate maps of putative chromosome structure among eight well assembled isolates, contigs were manually grouped based on nucmer alignments to mp102, or to mp053 for the mp102 strain. Only contigs >500 kb were evaluated. Of note, all strains except mp053 have a break in chromosome 1 at a large track (> 50 kb) of satellite sequence (satellite-7 in the repeat library) and are inferred to represent a single chromosome based on shared presence of satellite-7 at contig ends and alignments to this region in mp053.

The features of these accessory contigs additionally provided new insight into the biology of *M. phaseolina.* Many accessory contigs showed signatures of repeat-induced point mutations (RIP), a fungal genome defense mechanism that introduces cytosine to thymine transitions into repetitive DNA, which was not previously thought to be active in *M. phaseolina*^23^. Counter to the prevailing reports that *M. phaseolina* reproduces strictly through asexual propagation, the contrasting patterns of RIP across the twelve isolates suggest sexual reproduction has occurred at least once since their last common ancestor, as RIP only occurs immediately prior to sexual reproduction and meiosis (Table S26). In support of this hypothesis, we also observed two different mating type loci in isolates from both geographic regions suggestive of an intact bipolar heterothallic reproductive system (Table S27).

### *De novo* genome annotations

All final assemblies were annotated using a pipeline that integrated the output of multiple *de novo* gene prediction tools (Figure S14). Briefly, RepeatModeler v2.0.1 ^70^ using the Repbase database ^71^ was used to generate a custom repeat library that was combined with previously published transcriptome and annotated proteome data from *M. phaseolina*^23^ to generate initial gene predictions using MAKER2^72^. High quality predictions were used to train SNAP^73^, whose output was then combined with a *M. phaseolina*-specific Augustus parameter file^74^ and the output of self-trained GeneMark^75^ in a second round of MAKER2. High quality predictions were used to retrain parameter files for SNAP and Augustus, combined with the output of GeneMark, and used in a final round of MAKER2 to generate high-quality *de novo* gene annotations. To standardize annotations across all isolates, a crucial step for downstream comparative analyses, OrthoFiller was used for each genome to annotate any missing genes that were present in at least two other isolates^76^. All predicted proteins were annotated with Protein family (Pfam) domains and Gene Ontology (GO) terms using interproscan v5.48-83^77^. All assemblies were annotated for signatures of RIP using RIPper (default settings)^78^. To generate a custom repeat database for *M. phaseolina*, the output of default runs from each *M. phaseolina* genome were inspected manually to produce a fully classified and annotated library. The manual steps included removing redundancies and putative cellular genes, and ensuring repeats were properly annotated from beginning to end. Due to high divergence among individual repeat families (due to RIP), individual representative sequences were used in lieu of consensus sequences. Classification of most elements was determined using the standard schema provided by Wicker et al. ^26^. Non-canonical helitrons were discovered, and classified and annotated according to Chellapan et al. ^79^. Initially, repeats containing DUF3435 domains were annotated as *Cryptons*, and entered into the library. Many of these were later reclassified as *Starships* and given new names.

### Whole genome alignments and SNP tree

Whole genome alignments of the 12 new *M. phaseolina* genomes, a previously published *M. phaseolina* reference^23^ and an outgroup species *Botryosphaeria dothidea*^80^ were generated with Cactus ^81^ and projected against isolate mp102. MafFilter^82^ with recommended parameters^83^ was used to filter alignments and extract 3,227,097 SNP sites that were then concatenated and used to generate a maximum likelihood phylogeny with IQ-TREE v1.6.9 (-alrt 1000 -bb 1000 -bnni -m MFP+ASC)^84^. Total branch length distance between all pairs of isolates was calculated on the resulting phylogeny as a measurement of phylogenetic distance.

### Pangenome annotations and comparisons

All pairwise sequence comparisons between predicted proteins in the 12 *M. phaseolina* genomes were determined by diamond blastp (--more-sensitive -e 0.001 --max-hsps 1)^85^, and then used as input to the Pangloss pipeline(default settings)^86^ to determine one-to-one ortholog relationships between predicted genes using a combination of synteny- and sequence-based similarity metrics. In this way, each predicted gene from each isolate was assigned to an ortholog group, and each ortholog group had at most 1 representative sequence per isolate. We calculated pairwise percent identity in genome content between all isolates as the “Number of shared ortholog groups / Total number of non-redundant ortholog groups present across both genomes”. This process was then repeated for previously annotated genome datasets from six other fungal species in order to provide us with comparison points for evaluating *M. phaseolina*’s pangenome diversity (Table S5, Table S6). Datasets from the other species were chosen, when possible, to be composed of isolates from two distinct geographic locations, similar to the *M. phaseolina* dataset.

### Accessory gene hotspot analysis

We used BayesTraits v3.0.1 Multistate MCMC analysis^87^ in conjunction with the whole-genome SNP phylogeny to estimate the number of gains and losses experienced by each accessory ortholog group since the last common ancestor of the 12 *M. phaseolina* isolates. Each analysis was run for 10 million generations with a 2 million generation burn-in, sampling every 4,000 generations until 2,000 samples were collected. Because we lacked *a priori* expectations of parameter distributions, we seeded the means of all exponential priors from a uniform distribution ranging from 0-2 by using a hyper-prior, as recommended by the software authors. For each ortholog group, we derived the median probability of it being present or absent at each node in the phylogeny from the 2,000 samples of the posterior probability distribution. We then calculated the probability of gains and losses across the phylogeny by examining transitions between presence and absence states at each pair of parent and child nodes. To calculate the probability of gain of an ortholog group at a given child node, we multiplied the probability of its absence at the parent node by the probability of its presence at the child node. Similarly, to calculate the probability of loss of an ortholog group at a given child node, we multiplied the probability of its presence at the parent node by the probability of its absence at the child node. For example, if an ortholog group had a p(absent) = 0.75 at a parent node, and a p(present) = 0.75 at one of that parent node’s children, we calculated the p(gain) = 0.5625, and p(loss) = 0.0625 at the child node. We summed across all losses, and all gains in the phylogeny, to derive a phylogenetically- and probabilistically-weighted estimate of the number of gains and losses experienced by each ortholog group. Since the frequent gain and loss of TEs could obscure the gain and loss dynamics of host genes, we removed all ortholog groups where at least 1 member had either a predicted TE-associated protein domain or overlapped at least 50% with an annotated TE feature.

To identify regions of *M. phaseolina*’s genome where accessory ortholog group gains and losses were greater than expected by chance (i.e., hotspots), we derived a distribution based on the null expectation that ortholog groups are randomly distributed across the genome by sampling with replacement 5 million sets of 15 ortholog groups from randomly selected genomes. We calculated the total number of gains and losses per set, and found that <=1% of sets had a total number of gains and losses >=24.22. For each genome, we then calculated the total number of gains and losses in each 15 gene window on a 1-gene sliding window basis, and designated all windows >= 24.22 as hotspots, so that any hotspot had a <= 1% empirically-derived probability of being observed by chance. We then merged all overlapping hotspots into hotspot regions.

### Variant density

Genotyping structural mutations across a sample of individuals is challenging, especially on a genome-wide basis. We therefore developed a statistic analogous to Nei and Li’s average pairwise nucleotide diversity^88^, where we calculated the average length of different indel size classes in a given “focal” genome across all pairwise comparisons with other “query” genomes. We conducted all pairwise alignments of genome assemblies using minimap2 (-x asm10 -c --cs) and called SNP and indel variants using paftools(-l 1000 -L 5000)^89^. Using these alignments and variant calls, we classified all sequences in a focal genome into three types: sequence with no alignments (i.e., indels), sequence with 1:1 alignments (which included SNPs), and sequence with multiple alignments, for each pairwise comparison. We then overlaid this information with the coordinates of all hotspot regions for the focal genome, and calculated the average length of sequence impacted by SNPs and indels of various size classes (0.1-1kb, 1-10kb, >10kb) per 10kb of hotspot region. We then repeated this procedure for all background genomic regions. Since we were interested in aligning homologous regions, we only considered alignments to focal regions where <=5% sequence aligned to multiple locations in the query. To generate figures and calculate test statistics, we only considered focal regions that had valid alignments to 11 query genomes. Significant differences between the distributions of the average amount of sequence impacted by SNP and indels in hotspot and background regions were determined using the unpaired two sample Mann-Whitney U test implemented in the wilcox.test function in R.

### *Starship* search and classification

We initially discovered the *Voyager* element in *M. phaseolina* through progressively aligning the flanking sequences of large insertions to other genomes using MAFFT v7.407^90^ in order to identify “empty” insertion sites. We then used all predicted protein sequences within the *Voyager* reference element in *M. phaseolina* isolate mp102 (region ID: mp102|voyager|h11) as well as the SPOK3 sequence from *Enterprise* ^19^(GenBank accession: A0A516F180) as queries in a BLASTp search of the 12 *M. phaseolina* proteomes and 1274 published Ascomycete proteomes curated from public databases (BLAST filtering options: min. bit score = 50, max. Evalue = 1e^-4^; Table S12; gitlab.com/xonq/mycotools). Sequence similarity searches present a more comprehensive approach for mobile element annotation because they do not rely on the presence of polymorphic insertion sites. In *M. phaseolina*, all filtered hits to *Voyager* queries located within 100,000 bp of each other were merged into regions, and all regions with at least 4 hits were designated as *Starship*-like regions. Full length *Starship* regions were then characterized by progressively aligning flanking regions, as above. Regions whose insertion sites could not be resolved were bounded by genes at the 5’ and 3’ terminal ends. We aligned all regions to each other using nucmer and manually adjusted boundaries as necessary to include only sequence homologous to other regions. We repeated the above procedure for the larger genomic database, except we implemented a more conservative neighboring hit distance cutoff of 25,000 bp because in the vast majority of cases, we could not manually verify the region boundaries through pairwise alignments due to insufficient data. Genes within all regions were annotated with protein family (Pfam) domains using Interproscan v5.48-83^77^, and were additionally annotated with predicted functions and assigned to COG categories using eggNOG-mapper^91^.

### Phylogenetic classification of *M. phaseolina Starships*

All *Starships* and *Starship*-like regions in *M. phaseolina* were classified into distinct element types by comparing the phylogenetic relatedness of their DUF3435 captain sequences, following established approaches for classifying bacterial integrative and conjugative elements ^92^. Relationships were additionally supported using phylogenies of DUF3723, FRE, OCH1, and HET and NACHT domains from NLRs, and were used to identify the origin of degraded *Starship* regions. We first retrieved all hits to the DUF3435, DUF3723, FRE, and OCH1 sequences from the *Voyager* reference using a BLASTp search of the 12 *M. phaseolina* proteomes, and aligned all hits using MAFFT v7.407 (--auto)^90^. Due to the functional domain complexity of NLRs, we built individual phylogenetic trees for the HET and NACHT domains found most often in NLR genes (Table 2). We used hmmsearch to search the 12 *M. phaseolina* proteomes using profile HMMs (--incE 0.001), and aligned all hits to the query profile HMM using hmmalign ^93^. Alignments were trimmed using ClipKIT (default settings)^94^, and used as input to generate maximum likelihood phylogenies with SH-like approximate likelihood ratio test (SH-aLRT) and ultra fast bootstrap approximation (UFboot) support using IQ-TREE v2.0.5, where the best-fit substitution model was selected using ModelFinder (-m MFP -alrt 1000 -B 1000)^84, 95^.

### Tyrosine recombinase classification

An initial alignment of all DUF3435 sequences from the fully annotated starships (Table S14) was generated using MAFFT (defaults). This alignment was used as input to HHPRED ^96^ which identified the tyrosine recombinase domain of CRE recombinase (Protein Data Bank Accession: 1XO0) as the best hit to a conserved region of the alignment. A database of repeat elements with known tyrosine recombinase domains was collected from RepBase and HHPRED was used again to define the bounds of the tyrosine recombinase domains for each repeat family. This approach was then used for prokaryotic and plasmid elements from the ICEberg 2.0 database ^97^. For each sequence, all amino acids within the bounds of the tyrosine recombinase homology region were extracted, aligned using MAFFT v7.407 (--auto) ^90^, and trimmed as submitted to IQ-TREE as above. The tyrosine recombinase tree strongly supports the monophyly of the DUF3435 sequences in *Starship* elements in relation to other known tyrosine recombinase domains.

### Regression models

We built all phylogenetic generalized least squares regression models and assessed the significance of their parameter estimates using the “gls” function in ape v5.0 in conjunction with the SNP-based isolate tree (correlation = corBrownian, method = "ML")^98^. To test for the effect of *Voyager* on *M. phaseolina* growth, we used *Voyager* OCH1 copy number as the linear and quadratic predictor variables and either mean isolate AUDPC or AUGPC as the response variable for the pathogenicity and PDA experiments, respectively (Table S24).

### Statistical analyses and data visualization

All statistical analyses were conducted in R v4.1.1^99^. All Circos plots were generated using Circos^100^ and all other graphs were visualized using ggplot2^101^. All *Starship* schematics were either drawn manually or visualized using genoplotR^102^. All phylogenetic trees were visualized using ete3^103^.

## Supplemental Figure Legends

**Figure S1: Geographic origins and genomic characteristics of 12 newly sequenced *Macrophomina phaseolina* strains from North and South America.** A) The locations of soybean (*Glycine max*) fields in Paraguay and Ohio, U.S.A. from which individual strains were isolated from soil samples (Table macphIsolationData) B) *M. phaseolina* microsclerotia accumulation in a dissected soybean root, characteristic of charcoal rot disease. C) To the left, a maximum likelihood tree based on a 3,227,097 single nucleotide polymorphism alignment depicting relationships between all sequenced strains and the *Botryosphaeria dothidea* outgroup, with strain names highlighted according to their geographic origin. To the right of each branch tip is a bar chart summarizing the lengths in megabases of various genomic features in each draft assembly, followed by the assembly N50.

**Figure S2:. Chromosome maps depicting eight well assembled genomes of *M. phaseolina* isolates overlayed on the SNP-based phylogeny.** A) The consensus genome is inferred based on the chromosome structures of mp102 and mp053, the best assembled and most distantly related isolates. Chromosomes are colored based on inferred synteny as calculated from pairwise nuncmer alignments. Contigs are manually joined based on synteny to mp102, or to mp053 for the mp102 isolate. Contig breaks are depicted with a black vertical line (**|**), putative telomeres are shown as a gold diamond, *Starships* and *Starship*-like regions are colored dark gray. Thin, horizontal gray lines illustrate non-syntenic regions. Only contigs > 500 kb are illustrated, unplaced contigs are not illustrated. The phylogeny was edited from the maximum likelihood tree presented in Figure S1. B) Circos plot depicting pairwise alignments between mp102 and mp053. Chromosomes are colored as in A). Nucmer output was filtered to exclude alignments <14 kb, to avoid depicting conventional transposable element movement. Contigs and contig regions with no links are presumed to represent accessory genomic regions.

**Figure S3: Differences in *M. phaseolina*’s genome content accumulate rapidly and are concentrated in hotspot regions enriched for large structural polymorphisms.** A) A scatterplot depicting pairwise nucleotide- and gene-based identity as a function of phylogenetic distance between all non-redundant combinations of 12 *Macrophomina phaseolina* genomes. Predicted linear equations with shaded 95% confidence intervals derived from linear models are superimposed for each comparison type, along with p-value probabilities that the slopes do not differ significantly from 0. B) Rarefaction curves depicting the number of unique ortholog groups (OGs) recovered in subsamples of different genomic datasets, color-coded by fungal species consisting of *M. phaseolina* and several others included for comparison. Each point represents the mean of 1000 permutations, with error bars representing +/- standard deviation. To the right are estimates of Heaps Alpha, a measurement of pangenome openness, for each species. C) A Manhattan plot indicating the total number of gains and losses associated with all OGs in a given sliding window across the *M. phaseolina* mp102 genome. A dotted red line is drawn at the critical value used to identify hotspot regions, determined empirically using a randomly generated null distribution of expected gene gains and losses. D) Box-and-whisker plots summarizing the average amount of sequence per 10kb of a given focal region impacted by SNPs and indels of various sizes in hotspot and background regions, across all genomes. Significant differences between the distributions of sequence length in either region were determined by a one sided Mann Whitney U test.

**Figure S4: A maximum likelihood phylogeny of all 1890 DUF3435 homologs retrieved by BLASTp from the 12 *Macrophomina phaseolina* isolates and a database of 1274 published Ascomycete genomes using *Voyager*’s DUF3435 as a query.** Branches are color coded according to taxonomic class and sequences labeled by species. The phylogeny is rooted at a DUF3435 sequence retrieved from *Lentinula edodes* (Basidiomycetes). Sequences found in *Starships* or *Starship-*like regions are labeled by the region in which they are found, and captain sequences are additionally highlighted in red (Table S11, Table S13). Captain sequences are defined as the DUF3435 sequence located nearest to the 5’ terminal end of a *Starship* element and oriented in the 5’-3’ direction. Branch support labels indicate SH-aLRT and UFboot support, respectively. All branches with <80% SH-aLRT or <95% UFboot have been collapsed. A simplified, circularized version of this phylogeny is visualized in Figure 1A.

**Figure S5: A maximum likelihood phylogeny of all DUF3435 homologs found across 195 *Starships* and *Starship*-like regions from the 12 *Macrophomina phaseolina* isolates and a database of 1274 published Ascomycete genomes.** Branches are color coded according to taxonomic class and sequences labeled by species. The phylogeny is rooted at three DUF3435 sequences retrieved from Basidiomycetes. Branch support labels indicate SH-aLRT and UFboot support, respectively. All branches with <80% SH-aLRT or <95% UFboot have been collapsed. A simplified version of this phylogeny is visualized in Figure 1A. To the right of the tree is a heatmap matrix summarizing the number of predicted genes within each *Starship* and *Starship-*like region that are homologous to DUF3723, FRE, PLP and NLRs (Table S11, Table S13).

**Figure S6: A maximum likelihood phylogeny of all 165 DUF3435 homologs retrieved by BLASTp from the 12 *Macrophomina phaseolina* isolates using *Voyager*’s DUF3435 as a query, rooted at the last common ancestor of *Voyager*, *Argo*, *Defiant*, and *Phoenix* captain sequences.** Captain sequences are defined as the DUF3435 sequence located nearest to the 5’ terminal end of a *Starship* element and oriented in the 5’-3’ direction. Sequences found in *Starships* or *Starship-*like regions are labeled by the region in which they are found, and captain sequences are additionally highlighted in red (Table S11). Branch support labels indicate SH-aLRT and UFboot support, respectively. All branches with <80% SH-aLRT or <95% UFboot have been collapsed.

**Figure S7: A maximum likelihood phylogeny of the tyrosine recombinase domain from the DUF3435 captains of all fully annotated *Starships* presented in this study and reference sequences.** Sequences from characterized *Starships* are highlihgted in blue (Table S14). Tyrosine recombinase domains from known eukaryotic TEs, prokaryotic integrative elements, phages, and plasmids are included for reference. *Cryptons* are highlighted in green as they are mechanistically the most similar to the DUF34335 domain found in *Starships*, but are clearly distantly related and polyphyletic. The phylogeny is mid-point rooted. Branch support labels indicate SH-aLRT and UFboot support, respectively. All branches with <80% SH-aLRT and <95% UFboot have been collapsed.

**Figure S8: A maximum likelihood phylogeny of all 109 OCH1 homologs retrieved by BLASTp from the 12 *Macrophomina phaseolina* isolates using *Voyager*’s OCH1 as a query, all OCH1 sequences from *Botryosphaeria dothidea* strain LW-Hubei, and 5 reference sequences from functionally characterized genes in other fungi.** The phylogeny is midpoint rooted. Sequences found in *Starships* or *Starship-*like regions are labeled by the region in which they are found (Table S11). Branch support labels indicate SH-aLRT and UFboot support, respectively. All branches with <80% SH-aLRT and <95% UFboot have been collapsed.

**Figure S9: A maximum likelihood phylogeny of all 316 FRE homologs retrieved by BLASTp from the 12 *Macrophomina phaseolina* isolates using *Voyager*’s FRE as a query, and 16 reference sequences from functionally characterized genes in other fungi.** The phylogeny is mid-point rooted. Sequences found in *Starships* or *Starship-*like regions are labeled by the region in which they are found (Table S11). Branch support labels indicate SH-aLRT and UFboot support, respectively. All branches with <80% SH-aLRT and <95% UFboot have been collapsed.

**Figure S10: A maximum likelihood phylogeny of all 1007 predicted amino acid sequences containing HET domains (Protein family accession: PF06985) retrieved by HMMsearch from the 12 *Macrophomina phaseolina* isolates.** The phylogeny is mid-point rooted. Sequences found in *Starships* or *Starship-*like regions are labeled by the region in which they are found (Table S11). Branch support labels indicate SH-aLRT and UFboot support, respectively. All branches with <80% SH-aLRT and <95% UFboot have been collapsed.

**Figure S11: A maximum likelihood phylogeny of all 1441 predicted amino acid sequences containing NACHT domains (Protein family accession: PF05729) retrieved by HMMsearch from the 12 *Macrophomina phaseolina* isolates.** The phylogeny is mid-point rooted. Sequences found in *Starships* or *Starship-*like regions are labeled by the region in which they are found (Table S11). Branch support labels indicate SH-aLRT and UFboot support, respectively. All branches with <80% SH-aLRT and <95% UFboot have been collapsed.

**Figure S12: A maximum likelihood phylogeny of all 154 PLP homologs retrieved by BLASTp from the 12 *Macrophomina phaseolina* isolates using *Voyager*’s PLP as a query.** The phylogeny is mid-point rooted. Sequences found in *Starships* or *Starship-*like regions are labeled by the region in which they are found (Table S11). Branch support labels indicate SH-aLRT and UFboot support, respectively. All branches with <80% SH-aLRT and <95% UFboot have been collapsed.

**Figure S13: A maximum likelihood phylogeny of all 189 DUF3723 sequences retrieved by BLASTp from the 12 *Macrophomina phaseolina* isolates using *Voyager*’s DUF3723 as a query.** The phylogeny is mid-point rooted. Sequences found in *Starships* or *Starship-*like regions are labeled by the region in which they are found (Table S11). Branch support labels indicate SH-aLRT and UFboot support, respectively. All branches with <80% SH-aLRT and <95% UFboot have been collapsed.

**Figure S14: A flow chart describing the *de novo* annotation procedure of the 12 *M. phaseolina* genomes.** All steps indicating the use of particular software are visualized as rectangles with the software name indicated in bold; all output materials are visualized as parallelograms; all binary decision points are visualized as diamonds.

