## Supplemental Figures for "Giant *Starship* elements mobilize accessory genes in fungal genomes"

**Figure S1**

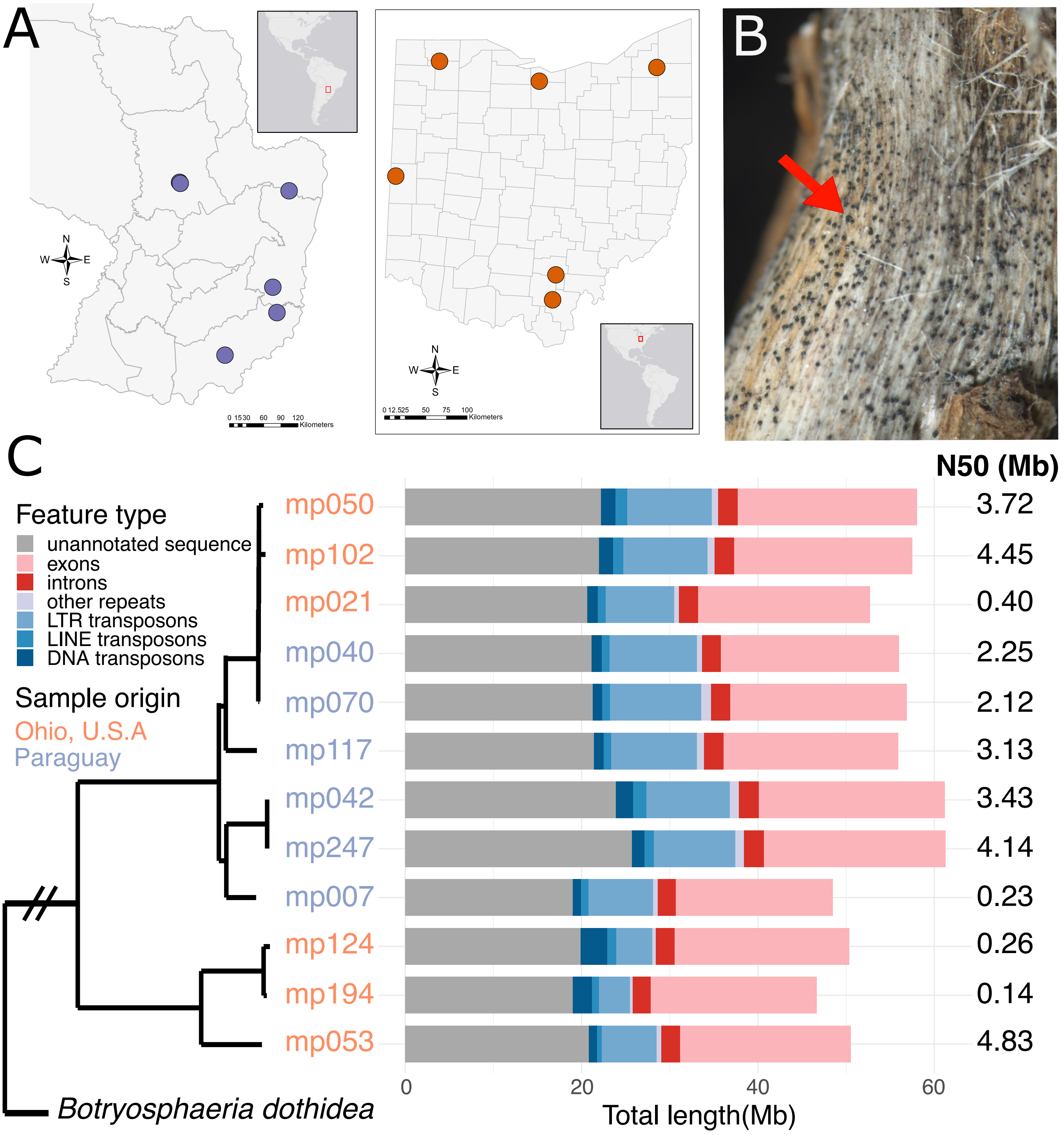
A
N
W
E
S
0 15 30 60 90 120
Kilometers
0 12.5 25 50 75 100
Kilometers
B
C
Feature type
unannotated sequence
exons
introns
other repeats
LTR transposons
LINE transposons
DNA transposons
Sample origin
Ohio, U.S.A
Paraguay
N50 (Mb)
mp050 3.72
mp102 4.45
mp021 0.40
mp040 2.25
mp070 2.12
mp117 3.13
mp042 3.43
mp247 4.14
mp007 0.23
mp124 0.26
mp194 0.14
mp053 4.83
Botryosphaeria dothidea
0
20
40
60
Total length(Mb)

### Figure S2

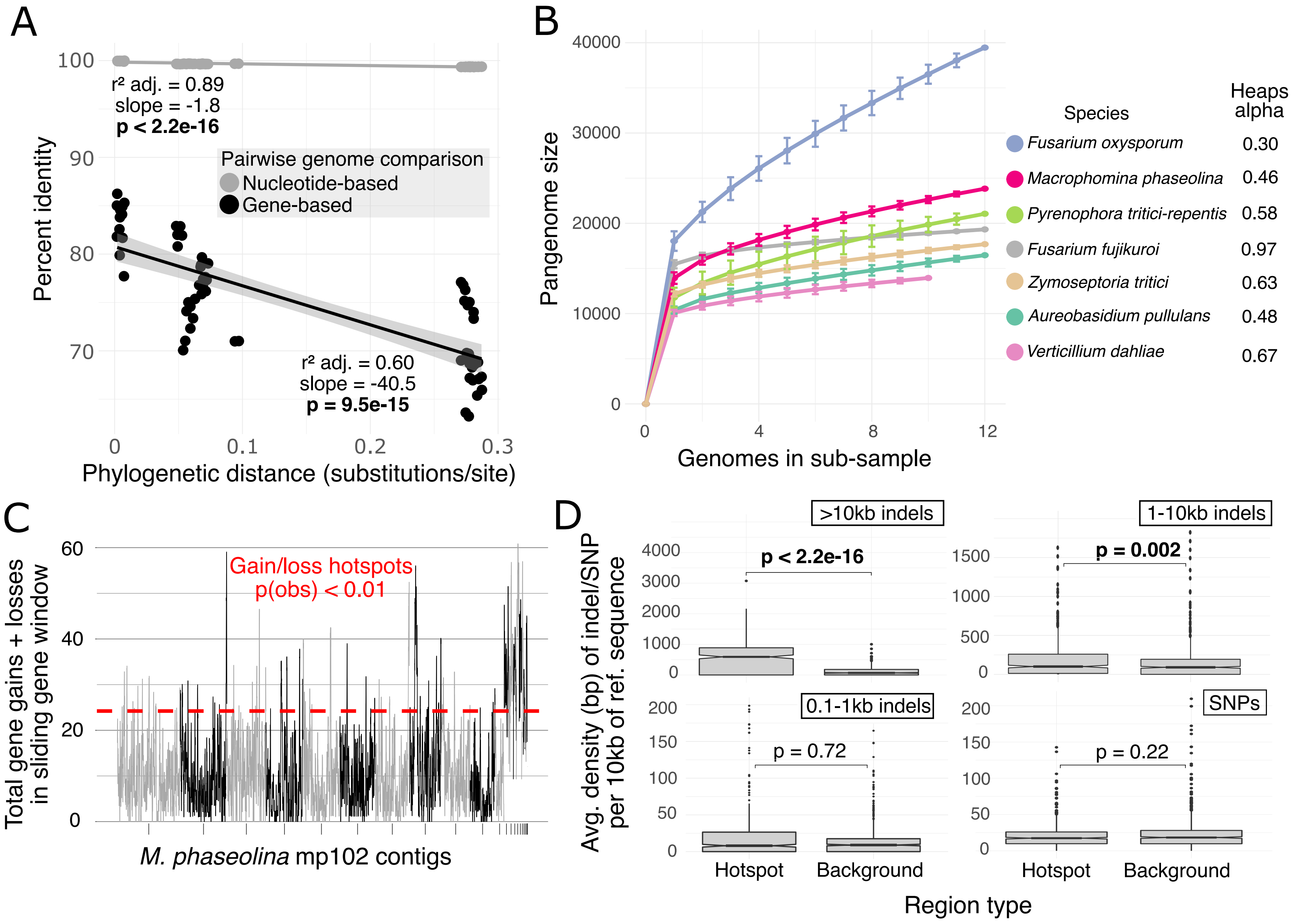

A
r² adj. = 0.89
slope = -1.8
p < 2.2e-16
Pairwise genome comparison
Nucleotide-based
Gene-based
r² adj. = 0.60
slope = -40.5
p = 9.5e-15
Percent identity
Phylogenetic distance (substitutions/site)
B
Pangenome size
Genomes in sub-sample
Species
Heaps alpha
Fusarium oxysporum 0.30
Macrophomina phaseolina 0.46
Pyrenophora tritici-repentis 0.58
Fusarium fujikuroi 0.97
Zymoseptoria tritici 0.63
Aureobasidium pullulans 0.48
Verticillium dahliae 0.67
C
Gain/loss hotspots
p(obs) < 0.01
Total gene gains + losses in sliding gene window
M. phaseolina mp102 contigs
D
>10kb indels
p < 2.2e-16
1-10kb indels
p = 0.002
0.1-1kb indels
p = 0.72
SNPs
p = 0.22
Avg. density (bp) of indel/SNP per 10kb of ref. sequence
Hotspot
Background
Region type

**Figure S3**

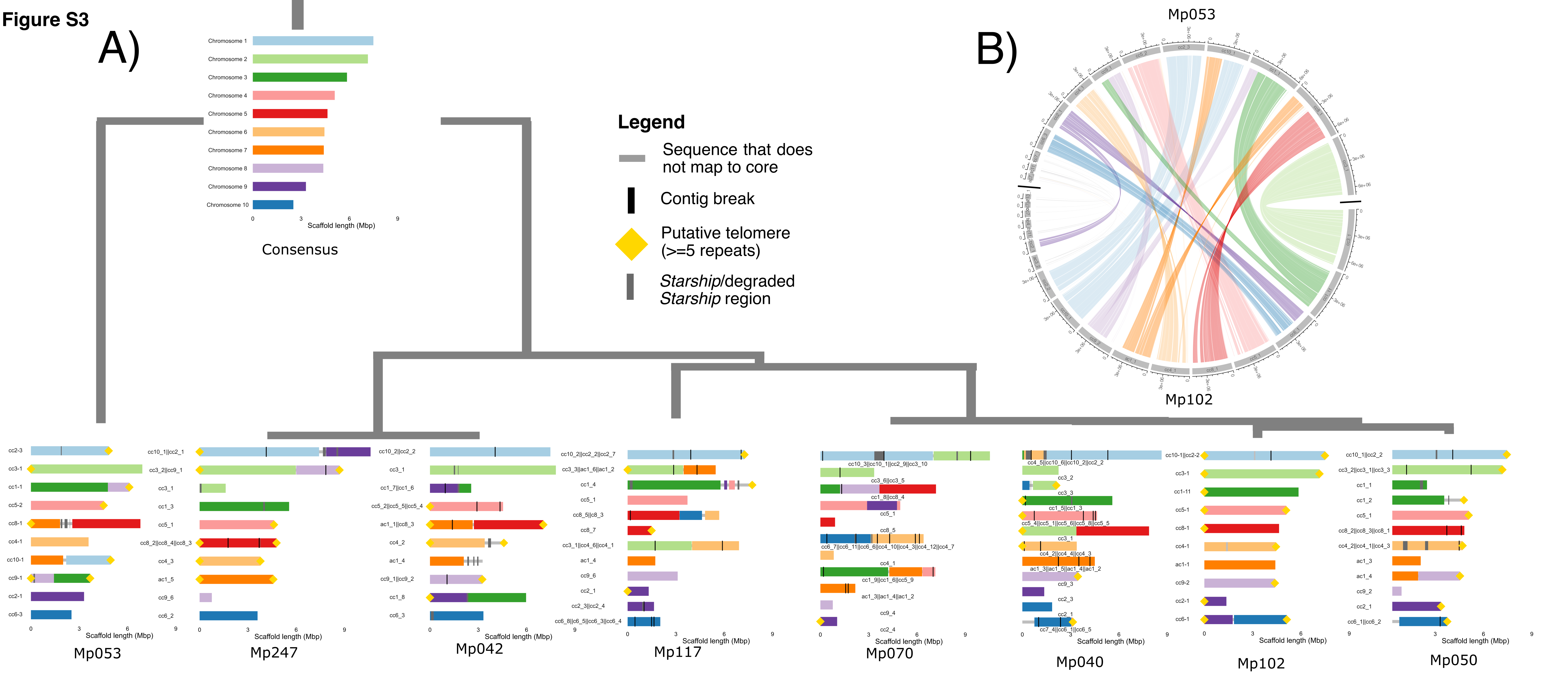
A)
B)
Chromosome 1
Chromosome 2
Chromosome 3
Chromosome 4
Chromosome 5
Chromosome 6
Chromosome 7
Chromosome 8
Chromosome 9
Chromosome 10
Scaffold length (Mbp)
Consensus
Legend
Sequence that does not map to core
Contig break
Putative telomere (>=5 repeats)
*Starship*/degraded *Starship* region
Mp053
Mp102
Mp053
Mp247
Mp042
Mp117
Mp070
Mp040
Mp102
Mp050

Figure S4

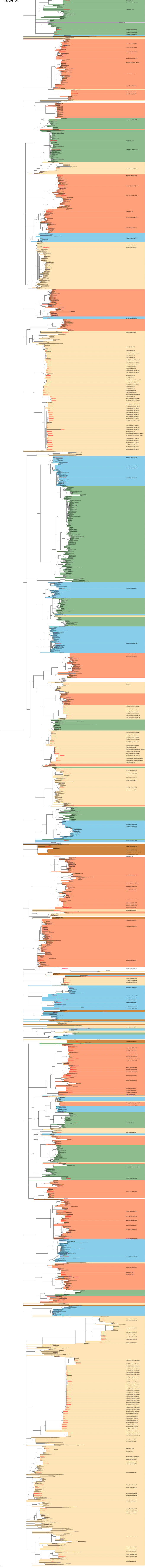

Figure S5

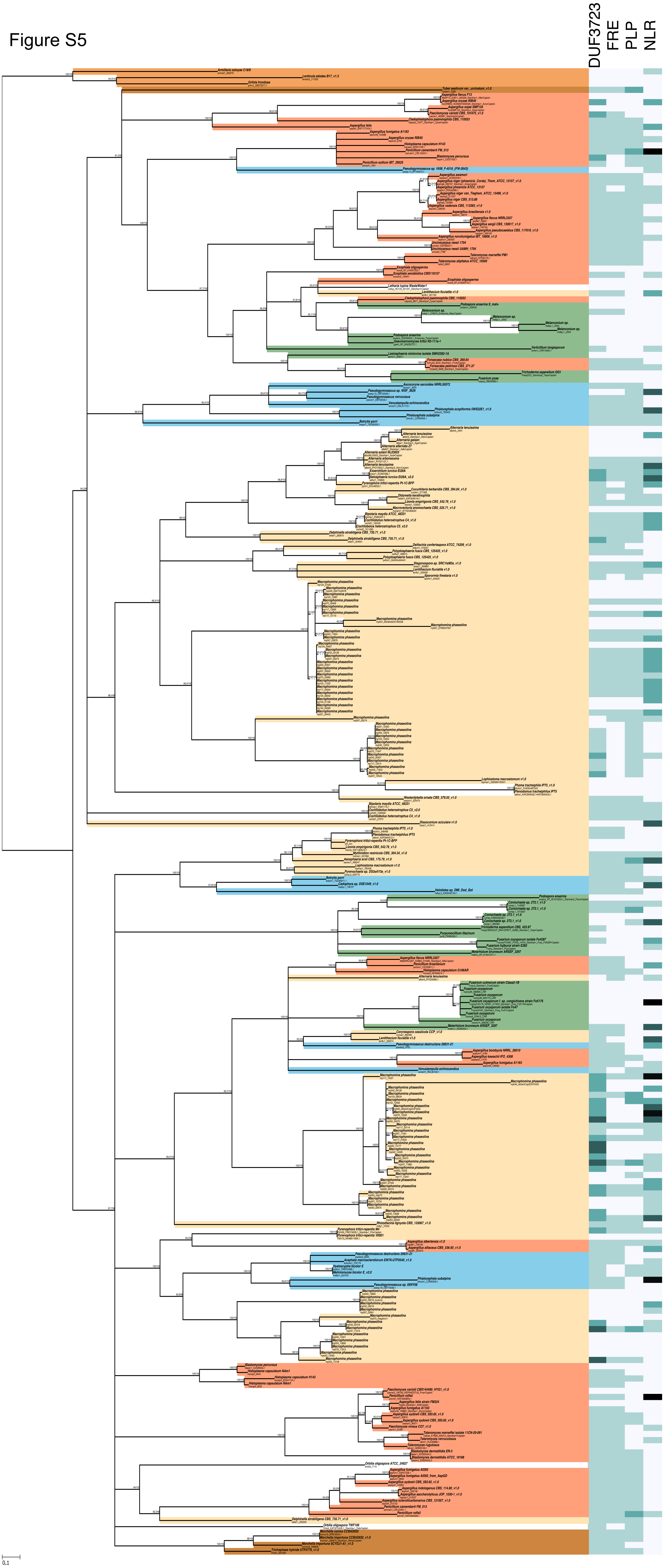

### Figure S6

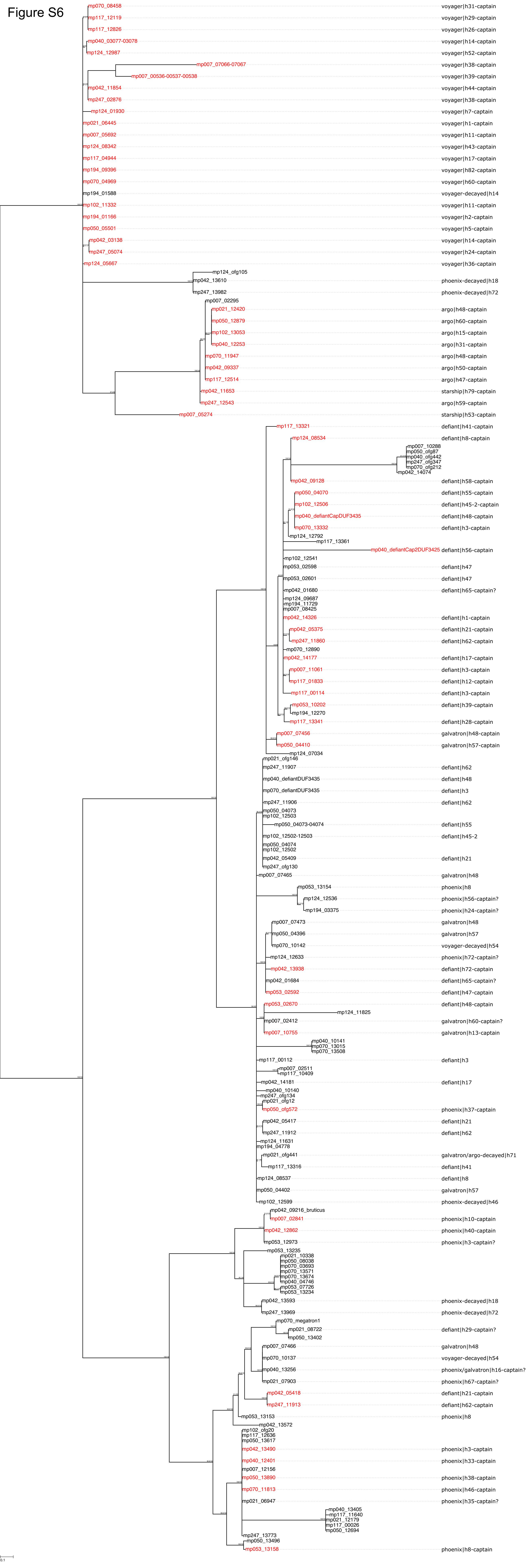

mp070_08458
voyager|h31-captain
mp117_12119
voyager|h29-captain
mp117_12826
voyager|h26-captain
mp040_03077-03078
voyager|h14-captain
mp124_12987
voyager|h52-captain
mp007_07066-07067
voyager|h38-captain
mp007_00536-00537-00538
voyager|h39-captain
mp042_11854
voyager|h44-captain
mp247_02876
voyager|h38-captain
mp124_01930
voyager|h7-captain
mp021_06445
voyager|h1-captain
mp007_05692
voyager|h11-captain
mp124_08342
voyager|h43-captain
mp117_04944
voyager|h17-captain
mp194_09396
voyager|h82-captain
mp070_04969
voyager|h60-captain
mp194_01588
voyager-decayed|h14
mp102_11332
voyager|h11-captain
mp194_01166
voyager|h2-captain
mp050_05501
voyager|h5-captain
mp042_03138
voyager|h14-captain
mp247_05074
voyager|h24-captain
mp124_05667
voyager|h36-captain
mp124_ofg105
mp042_13610
phoenix-decayed|h18
mp247_13982
phoenix-decayed|h72
mp007_02295
mp021_12420
argo|h48-captain
mp050_12879
argo|h60-captain
mp102_13053
argo|h15-captain
mp040_12253
argo|h31-captain
mp070_11947
argo|h48-captain
mp042_09337
argo|h50-captain
mp117_12514
argo|h47-captain
mp042_11653
starship|h79-captain
mp247_12543
argo|h59-captain
mp007_05274
starship|h53-captain
mp117_13321
defiant|h41-captain
mp124_08534
defiant|h8-captain
mp007_10288
mp050_ofg87
mp040_ofg442
mp247_ofg347
mp070_ofg212
mp042_14074
mp042_09128
defiant|h58-captain
mp050_04070
defiant|h55-captain
mp102_12506
defiant|h45-2-captain
mp040_defiantCapDUF3435
defiant|h48-captain
mp070_13332
defiant|h3-captain
mp124_12792
mp117_13361
mp040_defiantCap2DUF3425
defiant|h56-captain
mp102_12541
mp053_02598
defiant|h47
mp053_02601
defiant|h47
mp042_01680
defiant|h65-captain?
mp124_09687
mp194_11729
mp007_08425
mp042_14326
defiant|h1-captain
mp042_05375
defiant|h21-captain
mp247_11860
defiant|h62-captain
mp070_12890
mp042_14177
defiant|h17-captain
mp007_11061
defiant|h3-captain
mp117_01833
defiant|h12-captain
mp117_00114
defiant|h3-captain
mp053_10202
defiant|h39-captain
mp194_12270
mp117_13341
defiant|h28-captain
mp007_07456
galvatron|h48-captain
mp050_04410
galvatron|h57-captain
mp124_07034
mp021_ofg146
mp247_11907
defiant|h62
mp040_defiantDUF3435
defiant|h48
mp070_defiantDUF3435
defiant|h3
mp247_11906
defiant|h62
mp050_04073
mp102_12503
mp050_04073-04074
defiant|h55
mp102_12502-12503
defiant|h45-2
mp050_04074
mp102_12502
mp042_05409
defiant|h21
mp247_ofg130
mp007_07465
galvatron|h48
mp053_13154
phoenix|h8
mp124_12536
phoenix|h56-captain?
mp194_03375
phoenix|h24-captain?
mp007_07473
galvatron|h48
mp050_04396
galvatron|h57
mp070_10142
voyager-decayed|h54
mp124_12633
phoenix|h72-captain?
mp042_13938
defiant|h72-captain
mp042_01684
defiant|h65-captain?
mp053_02592
defiant|h47-captain
mp053_02670
defiant|h48-captain
mp124_11825
mp007_02412
galvatron|h60-captain?
mp007_10755
galvatron|h13-captain
mp040_10141
mp070_13015
mp070_13508
mp117_00112
defiant|h3
mp007_02511
mp117_10409
mp042_14181
defiant|h17
mp040_10140
mp247_ofg134
mp021_ofg12
mp050_ofg572
phoenix|h37-captain
mp042_05417
defiant|h21
mp247_11912
defiant|h62
mp124_11631
mp194_04778
mp021_ofg441
galvatron/argo-decayed|h71
mp117_13316
defiant|h41
mp124_08537
defiant|h8
mp050_04402
galvatron|h57
mp102_12599
phoenix-decayed|h46
mp042_09216_bruticus
mp007_02841
phoenix|h10-captain
mp042_12862
phoenix|h40-captain
mp053_12973
phoenix|h3-captain?
mp053_13235
mp021_10338
mp050_08038
mp070_03693
mp070_13571
mp070_13674
mp040_04746
mp053_07726
mp053_13234
mp042_13593
phoenix-decayed|h18
mp247_13969
phoenix-decayed|h72
mp070_megatron1
mp021_08722
defiant|h29-captain?
mp050_13402
mp007_07466
galvatron|h48
mp070_10137
voyager-decayed|h54
mp040_13256
phoenix/galvatron|h16-captain?
mp021_07903
phoenix|h67-captain?
mp042_05418
defiant|h21-captain
mp247_11913
defiant|h62-captain
mp053_13153
phoenix|h8
mp042_13572
mp102_ofg20
mp117_12636
mp050_13617
mp042_13490
phoenix|h3-captain
mp040_12401
phoenix|h33-captain
mp007_12156
mp050_13890
phoenix|h38-captain
mp070_11813
phoenix|h46-captain
mp021_06947
phoenix|h35-captain?
mp040_13405
mp117_11640
mp021_12179
mp117_00026
mp050_12694
mp247_13773
mp050_13496
mp053_13158
phoenix|h8-captain
0.1

Figure S7

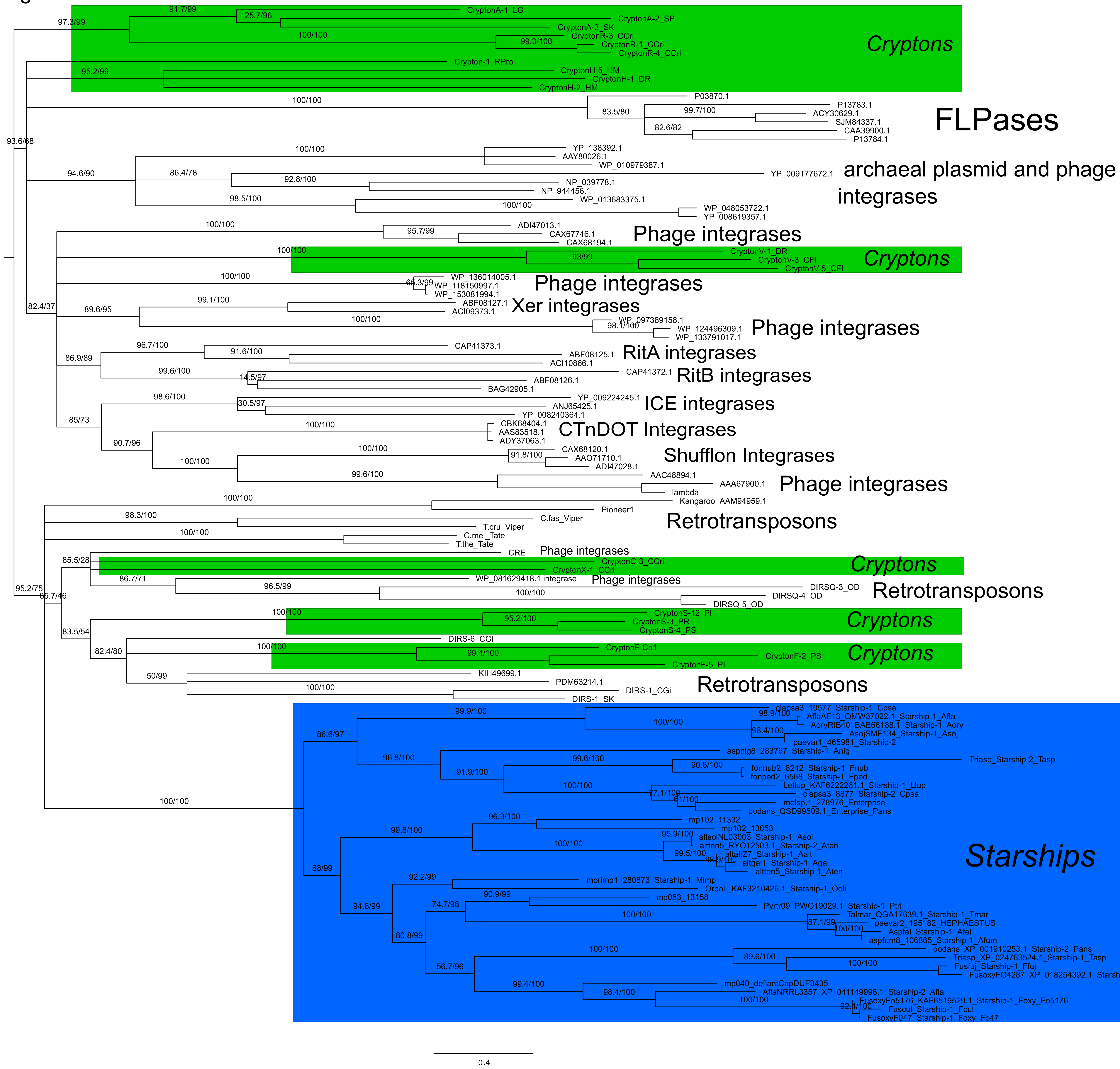

CryptonA-1_LG
CryptonA-2_SP
CryptonA-3_SK
CryptonR-3_CCri
CryptonR-1_CCri
CryptonR-4_CCri
Crypton-1_RPro
CryptonH-5_HM
CryptonH-1_DR
CryptonH-2_HM
Cryptons
P03870.1
P13783.1
ACY30629.1
SJM84337.1
CAA39900.1
P13784.1
FLPases
YP_138392.1
AAY80026.1
WP_010979387.1
YP_009177672.1
NP_039778.1
NP_944456.1
WP_013683375.1
WP_048053722.1
YP_008619357.1
archaeal plasmid and phage integrases
ADI47013.1
CAX67746.1
CAX68194.1
Phage integrases
CryptonV-1_DR
CryptonV-3_CFI
CryptonV-5_CFI
Cryptons
WP_136014005.1
WP_118150997.1
WP_153081994.1
Phage integrases
ABF08127.1
ACI09373.1
Xer integrases
WP_097389158.1
WP_124496309.1
WP_133791017.1
Phage integrases
CAP41373.1
ABF08125.1
ACI10866.1
RitA integrases
CAP41372.1
ABF08126.1
BAG42905.1
RitB integrases
YP_009224245.1
ANJ65425.1
YP_008240364.1
ICE integrases
CBK68404.1
AAS83518.1
ADY37063.1
CTnDOT Integrases
CAX68120.1
AAO71710.1
ADI47028.1
Shufflon Integrases
AAC48894.1
AAA67900.1
lambda
Kangaroo_AAM94959.1
Phage integrases
Pioneer1
C.fas_Viper
T.cru_Viper
C.mel_Tate
T.the_Tate
Retrotransposons
CRE
Phage integrases
CryptonC-3_CCri
CryptonX-1_CCri
Cryptons
WP_081629418.1 integrase
Phage integrases
DIRSQ-3_OD
DIRSQ-4_OD
DIRSQ-5_OD
Retrotransposons
CryptonS-12_PI
CryptonS-3_PR
CryptonS-4_PS
Cryptons
DIRS-6_CGi
CryptonF-Cn1
CryptonF-2_PS
CryptonF-5_PI
Cryptons
KIH49699.1
PDM63214.1
DIRS-1_CGi
DIRS-1_SK
Retrotransposons
clapsa3_10577_Starship-1_Cpsa
AflaAF13_QMW37022.1_Starship-1_Afla
AoryRIB40_BAE66188.1_Starship-1_Aory
AsojSMF134_Starship-1_Asoj
paevar1_465981_Starship-2
aspnig8_283767_Starship-1_Anig
Triasp_Starship-2_Tasp
fonnub2_8242_Starship-1_Fnub
fonped2_6568_Starship-1_Fped
Letlup_KAF6222261.1_Starship-1_Llup
clapsa3_8877_Starship-2_Cpsa
melsp.1_278976_Enterprise
podans_QSD99509.1_Enterprise_Pans
mp102_11332
mp102_13053
altsolNL03003_Starship-1_Asol
alten5_RYO12503.1_Starship-2_Aten
altaltZ7_Starship-1_Aalt
altgai1_Starship-1_Agai
alten5_Starship-1_Aten
morimp1_280873_Starship-1_Mimp
Orboli_KAF3210426.1_Starship-1_Ooli
mp053_13158
Pyrtr09_PWO19029.1_Starship-1_Ptri
Talmar_QGA17639.1_Starship-1_Tmar
paevar2_195182_HEPHAESTUS
Aspfel_Starship-1_Afel
aspfum8_106865_Starship-1_Afum
podans_XP_001910253.1_Starship-2_Pans
Triasp_XP_024763524.1_Starship-1_Tasp
Fusfuj_Starship-1_Ffuj
FusoxyFO4287_XP_018254392.1_Starsh
mp040_defiantCapDUF3435
AflaNRRL3357_XP_041149996.1_Starship-2_Afla
FusoxyFo5176_KAF6519529.1_Starship-1_Foxy_Fo5176
Fuscul_Starship-1_Fcul
FusoxyF047_Starship-1_Foxy_Fo47
Starships
0.4

Figure S8

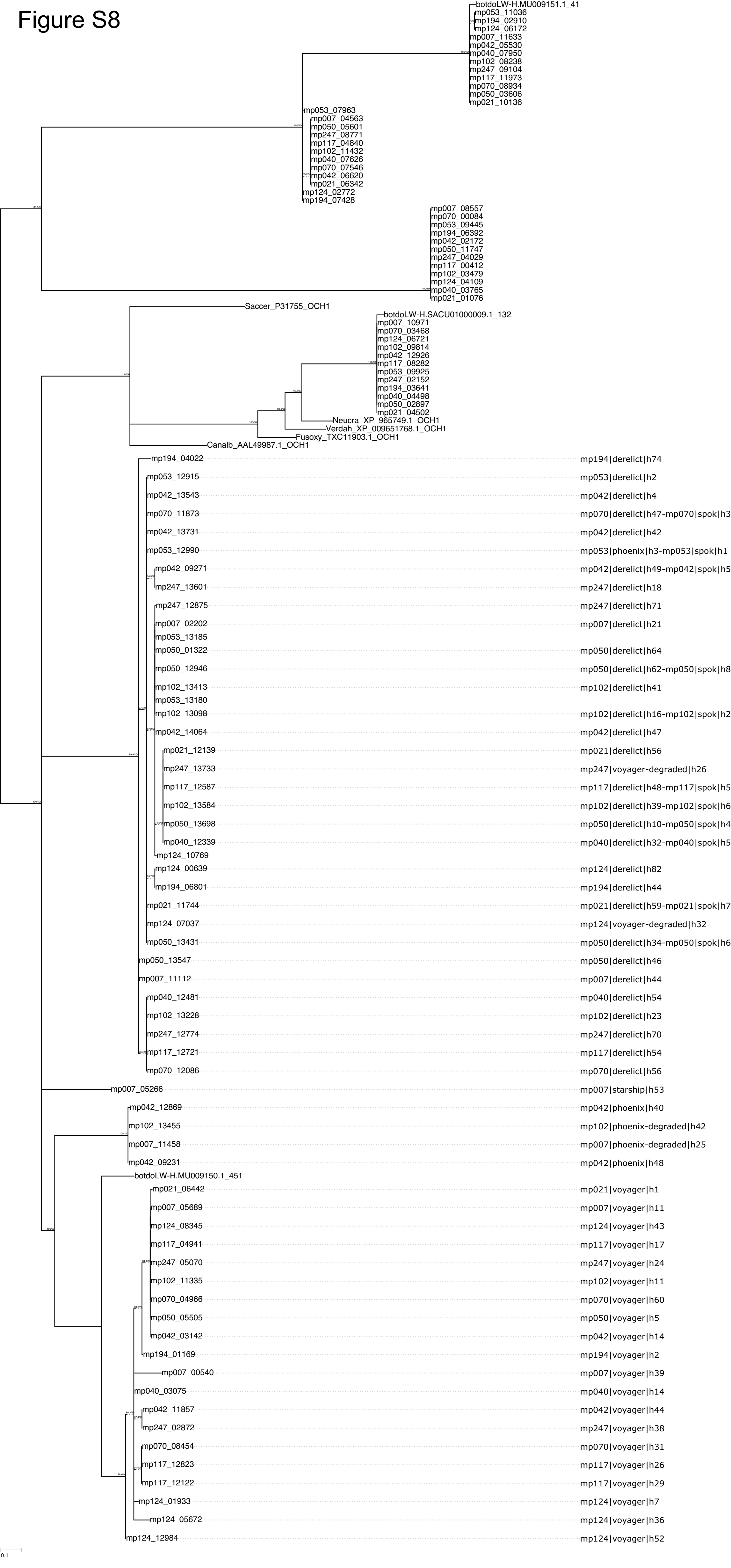

Figure S9

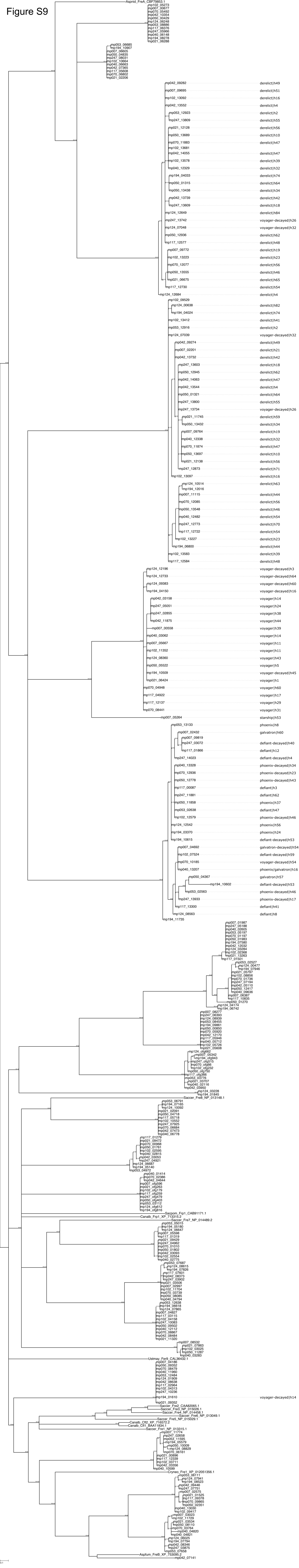

### Figure S10

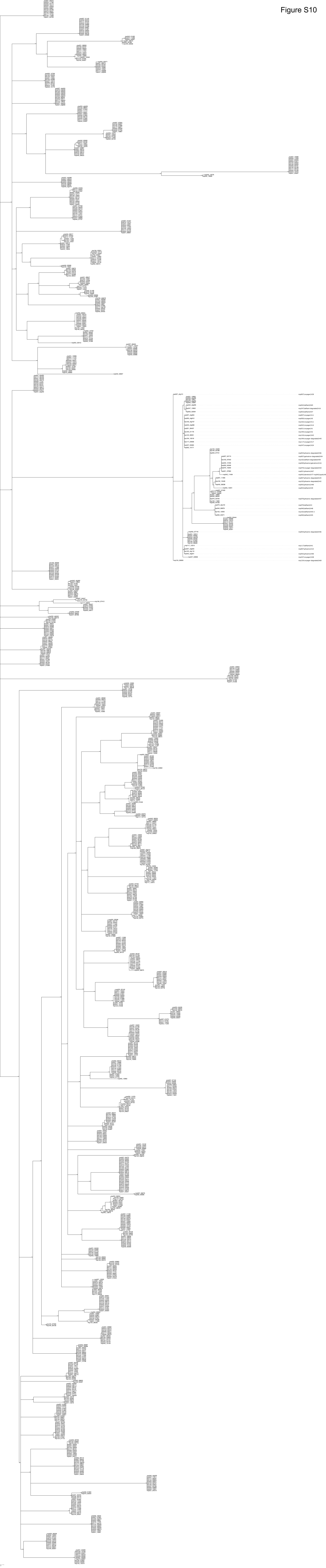

### Figure S11

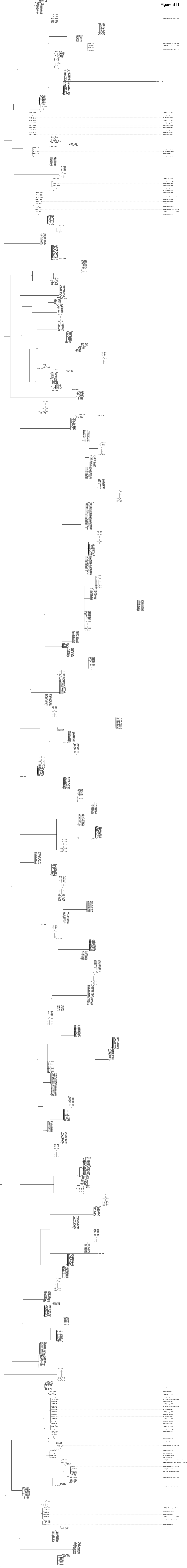

### Figure S12

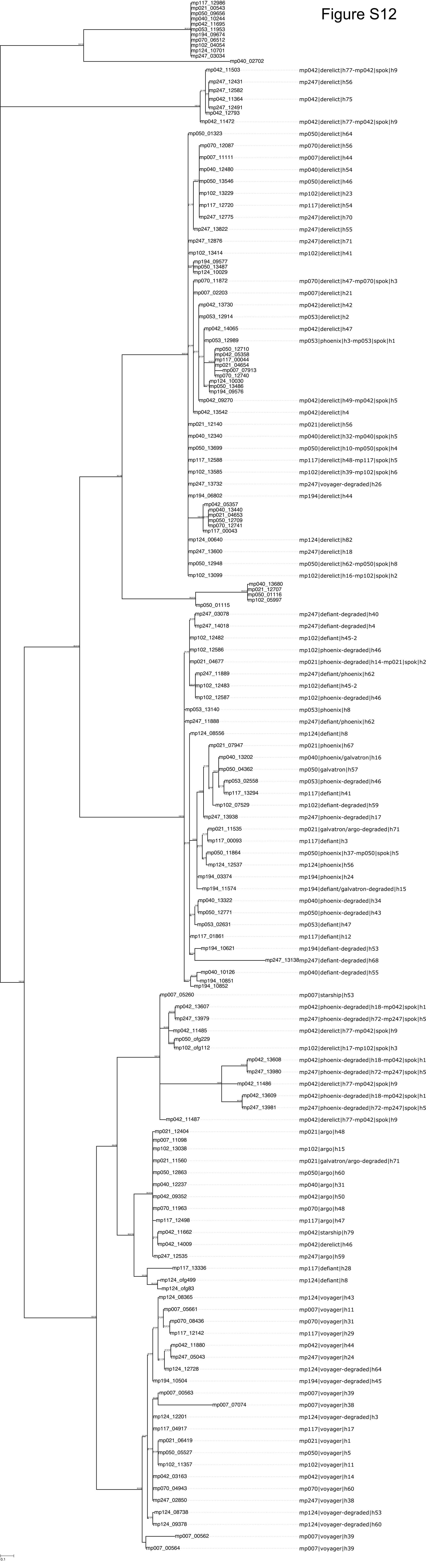

mp117_12986
mp021_00543
mp050_09656
mp040_10244
mp042_11695
mp053_11953
mp194_09674
mp070_06512
mp102_04054
mp124_10701
mp247_03034
mp040_02702
mp042_11503 mp042|derelict|h77-mp042|spok|h9
mp247_12431 mp247|derelict|h56
mp247_12582
mp042_11364 mp042|derelict|h75
mp247_12491
mp042_12793
mp042_11472 mp042|derelict|h77-mp042|spok|h9
mp050_01323 mp050|derelict|h64
mp070_12087 mp070|derelict|h56
mp007_11111 mp007|derelict|h44
mp040_12480 mp040|derelict|h54
mp050_13546 mp050|derelict|h46
mp102_13229 mp102|derelict|h23
mp117_12720 mp117|derelict|h54
mp247_12775 mp247|derelict|h70
mp247_13822 mp247|derelict|h55
mp247_12876 mp247|derelict|h71
mp102_13414 mp102|derelict|h41
mp194_09577
mp050_13487
mp124_10029
mp070_11872 mp070|derelict|h47-mp070|spok|h3
mp007_02203 mp007|derelict|h21
mp042_13730 mp042|derelict|h42
mp053_12914 mp053|derelict|h2
mp042_14065 mp042|derelict|h47
mp053_12989 mp053|phoenix|h3-mp053|spok|h1
mp050_12710
mp042_05358
mp117_00044
mp021_04654
mp007_07913
mp070_12740
mp124_10030
mp050_13486
mp194_09576
mp042_09270 mp042|derelict|h49-mp042|spok|h5
mp042_13542 mp042|derelict|h4
mp021_12140 mp021|derelict|h56
mp040_12340 mp040|derelict|h32-mp040|spok|h5
mp050_13699 mp050|derelict|h10-mp050|spok|h4
mp117_12588 mp117|derelict|h48-mp117|spok|h5
mp102_13585 mp102|derelict|h39-mp102|spok|h6
mp247_13732 mp247|voyager-degraded|h26
mp194_06802 mp194|derelict|h44
mp042_05357
mp040_13440
mp021_04653
mp050_12709
mp070_12741
mp117_00043
mp124_00640 mp124|derelict|h82
mp247_13600 mp247|derelict|h18
mp050_12948 mp050|derelict|h62-mp050|spok|h8
mp102_13099 mp102|derelict|h16-mp102|spok|h2
mp040_13680
mp021_12707
mp050_01116
mp102_05997
mp050_01115
mp247_03078 mp247|defiant-degraded|h40
mp247_14018 mp247|defiant-degraded|h4
mp102_12482 mp102|defiant|h45-2
mp102_12586 mp102|phoenix-degraded|h46
mp021_04677 mp021|phoenix-degraded|h14-mp021|spok|h2
mp247_11889 mp247|defiant/phoenix|h62
mp102_12483 mp102|defiant|h45-2
mp102_12587 mp102|phoenix-degraded|h46
mp053_13140 mp053|phoenix|h8
mp247_11888 mp247|defiant/phoenix|h62
mp124_08556 mp124|defiant|h8
mp021_07947 mp021|phoenix|h67
mp040_13202 mp040|phoenix/galvatron|h16
mp050_04362 mp050|galvatron|h57
mp053_02558 mp053|phoenix-degraded|h46
mp117_13294 mp117|defiant|h41
mp102_07529 mp102|defiant-degraded|h59
mp247_13938 mp247|phoenix-degraded|h17
mp021_11535 mp021|galvatron/argo-degraded|h71
mp117_00093 mp117|defiant|h3
mp050_11864 mp050|phoenix|h37-mp050|spok|h5
mp124_12537 mp124|phoenix|h56
mp194_03374 mp194|phoenix|h24
mp194_11574 mp194|defiant/galvatron-degraded|h15
mp040_13322 mp040|phoenix-degraded|h34
mp050_12771 mp050|phoenix-degraded|h43
mp053_02631 mp053|defiant|h47
mp117_01861 mp117|defiant|h12
mp194_10621 mp194|defiant-degraded|h53
mp247_13138 mp247|defiant-degraded|h68
mp040_10126 mp040|defiant-degraded|h55
mp194_10851
mp194_10852
mp007_05260 mp007|starship|h53
mp042_13607 mp042|phoenix-degraded|h18-mp042|spok|h1
mp247_13979 mp247|phoenix-degraded|h72-mp247|spok|h5
mp042_11485 mp042|derelict|h77-mp042|spok|h9
mp050_ofg229
mp102_ofg112 mp102|derelict|h17-mp102|spok|h3
mp042_13608 mp042|phoenix-degraded|h18-mp042|spok|h1
mp247_13980 mp247|phoenix-degraded|h72-mp247|spok|h5
mp042_11486 mp042|derelict|h77-mp042|spok|h9
mp042_13609 mp042|phoenix-degraded|h18-mp042|spok|h1
mp247_13981 mp247|phoenix-degraded|h72-mp247|spok|h5
mp042_11487 mp042|derelict|h77-mp042|spok|h9
mp021_12404 mp021|argo|h48
mp007_11098
mp102_13038 mp102|argo|h15
mp021_11560 mp021|galvatron/argo-degraded|h71
mp050_12863 mp050|argo|h60
mp040_12237 mp040|argo|h31
mp042_09352 mp042|argo|h50
mp070_11963 mp070|argo|h48
mp117_12498 mp117|argo|h47
mp042_11662 mp042|starship|h79
mp042_14009 mp042|derelict|h46
mp247_12535 mp247|argo|h59
mp117_13336 mp117|defiant|h28
mp124_ofg499 mp124|defiant|h8
mp124_ofg83
mp124_08365 mp124|voyager|h43
mp007_05661 mp007|voyager|h11
mp070_08436 mp070|voyager|h31
mp117_12142 mp117|voyager|h29
mp042_11880 mp042|voyager|h44
mp247_05043 mp247|voyager|h24
mp124_12728 mp124|voyager-degraded|h64
mp194_10504 mp194|voyager-degraded|h45
mp007_00563 mp007|voyager|h39
mp007_07074 mp007|voyager|h38
mp124_12201 mp124|voyager-degraded|h3
mp117_04917 mp117|voyager|h17
mp021_06419 mp021|voyager|h1
mp050_05527 mp050|voyager|h5
mp102_11357 mp102|voyager|h11
mp042_03163 mp042|voyager|h14
mp070_04943 mp070|voyager|h60
mp247_02850 mp247|voyager|h38
mp124_08738 mp124|voyager-degraded|h53
mp124_09378 mp124|voyager-degraded|h60
mp007_00562 mp007|voyager|h39
mp007_00564 mp007|voyager|h39
0.1

Figure S13

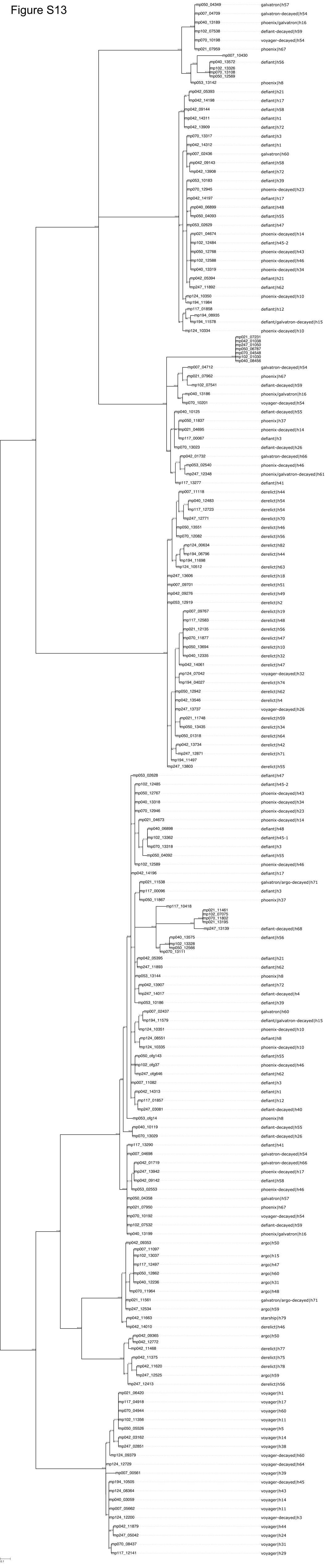

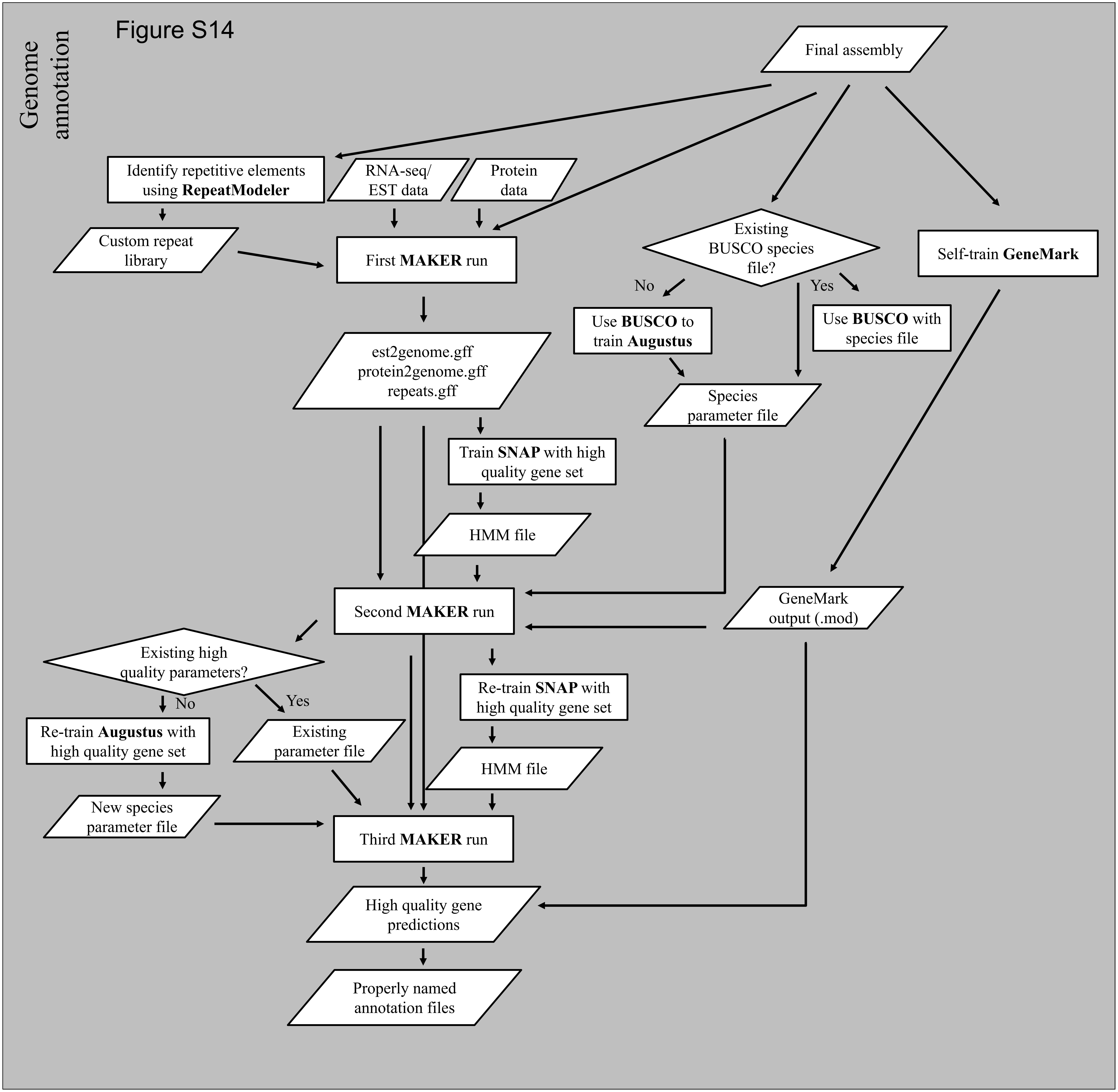
Figure S14
Genome annotation
Final assembly
Identify repetitive elements using **RepeatModeler**
RNA-seq/ EST data
Protein data
Custom repeat library
First **MAKER** run
Existing BUSCO species file?
No
Yes
Use **BUSCO** to train **Augustus**
Use **BUSCO** with species file
Self-train **GeneMark**
est2genome.gff
protein2genome.gff
repeats.gff
Species parameter file
Train **SNAP** with high quality gene set
HMM file
Second **MAKER** run
GeneMark output (.mod)
Existing high quality parameters?
No
Yes
Re-train **SNAP** with high quality gene set
Re-train **Augustus** with high quality gene set
Existing parameter file
HMM file
New species parameter file
Third **MAKER** run
High quality gene predictions
Properly named annotation files
